# ENOX2 NADH Oxidase: A BCR-ABL1-dependent cell surface and secreted redox protein in chronic myeloid leukemia (CML)

**DOI:** 10.1101/2021.01.23.427819

**Authors:** Seda Baykal-Köse, Maud Voldoire, Christophe Desterke, Nathalie Sorel, Emilie Cayssials, Hyacinthe Johnson-Ansah, Agnes Guerci-Bresler, Annelise Bennaceur-Griscelli, Jean-Claude Chomel, Ali G Turhan

## Abstract

Chronic myeloid leukemia (CML) is a myeloproliferative neoplasm caused by the acquisition of *BCR-ABL1* fusion in a hematopoietic stem cell. We identified the *ENOX2* gene as up-regulated in *BCR-ABL1*-expressing UT-7 cell lines through a transcriptome assay. The oncofoetal ENOX2 protein (Ecto-Nicotinamide Adenine Dinucleotide Oxidase Disulfide Thiol Exchanger 2) is expressed on the external plasma membrane surface of cancer cells and can be released in cancer patients’ serum. Considering these data, we studied ENOX2 expression in CML cell lines and patients using quantitative RT-PCR, western-blots, the ELISA method, and transcriptomic dataset reanalysis. We confirmed increased *ENOX2* mRNA expression in the *BCR-ABL1*-expressing UT-7 cell line. Comparable results were obtained in CML patients at diagnosis. Western-blot analyses on UT-7 and TET-inducible Ba/F3 cell lines established the up-regulation of ENOX2 protein. BCR-ABL1 has been found to induce ENOX2 overexpression in a kinase-dependent manner. In a series of 41 patients with CML, ELISA assays showed a highly significant increase of ENOX2 protein levels in the plasma of patients with CML (p < 0.0001) as compared to controls (n=28). Transcriptomic dataset (GSE4170) reanalyzes have shown specific *ENOX2* mRNA overexpression in the chronic phase of the disease. Bioinformatic analyses identified several genes whose mRNA expression was positively correlated to *ENOX2*. Some of them encode proteins involved in cellular functions compatible with the growth deregulation observed in CML. All in all, our results demonstrate for the first time the upregulation of a secreted Redox protein in a BCR-ABL1-dependent manner in CML. Our data suggest that ENOX2 (through its transcriptional program) plays a significant role in the BCR-ABL1 leukemogenesis. Further studies are required to clarify the relationship between BCR-ABL1 and ENOX2.

## Introduction

Chronic myeloid leukemia (CML) is a myeloproliferative neoplasm characterized by the *BCR-ABL1* molecular rearrangement resulting from t(9;22)(q34;q11) translocation. The natural history of the disease has been radically modified by tyrosine kinase inhibitors (TKIs) of first-(Imatinib), second-(Dasatinib, Nilotinib, Bosutinib), and third-generation (Ponatinib). Nowadays, the life expectancy of patients diagnosed in chronic phase-CML (CP-CML) and responding to TKI therapy is very similar to that of the general population [1,2]. In most cases, the current management of CP-CML patients prevents a progression to accelerated phase or blast crisis [3]. Among hematological malignancies, CML provides a unique biological model to apprehend leukemogenesis, genetic instability, targeted therapy management, drug resistance, or stem cell persistence [4]. The constitutively active BCR-ABL1 tyrosine kinase oncoprotein is directly responsible for the oncogenic process. It initiates uncontrolled granulocyte proliferation, decreased adhesion to bone marrow niche cells, apoptosis inhibition, and increased genetic instability. Most of the aberrant signaling pathways downstream BCR-ABL1 (particularly well-known MAPK, PI3K, MYC, STAT pathways) have been known for a long time [5].

Using the human GM-CSF-dependent erythro/megakaryoblastic UT-7 cell line, our group previously uncovered novel non-conventional actors downstream to the BCR-ABL1 signaling [6–8]. In the present work, we highlight *ENOX2* (Ecto-Nicotinamide Adenine Dinucleotide Oxidase Disulfide Thiol Exchanger 2) as up-regulated in *BCR-ABL1*-positive cell lines. ENOX2, also named APK1 antigen, COVA1 (Cytosolic Ovarian Carcinoma Antigen 1), or tNOX (Tumor-Associated Nicotinamide Adenine Dinucleotide Oxidase), is a growth-related cell surface protein. ENOX2 dimers combine two main enzymatic oscillatory activities (hydroquinone NADH oxidase and protein disulfide-thiol oxidoreductase) that alternate with a period length of approximately 22 minutes, generating an ultradian cellular biological clock of 22 hours (S1 Fig) [9–11].

Increased *ENOX2* mRNA expression was also found in primary cells from CP-CML patients at diagnosis. Based on western-blot experiments carried out on UT-7 and inducible Ba/F3 cell lines, we confirmed ENOX2 up-regulation at the protein level, and we demonstrated that this phenomenon was linked to the BCR-ABL1 tyrosine kinase activity. In addition, a significant increase of ENOX2 protein levels was observed in the plasma of patients at the time of diagnosis. Reanalyzing a publicly available database, we found that *ENOX2* mRNA expression was characteristic of the chronic phase of CML. Moreover, several genes’ mRNA expression was shown to be positively correlated with *ENOX2*. Some of these genes encode proteins involved in cellular functions compatible with the dysregulated hematopoiesis observed in CP-CML.

## Materials and Methods

### UT-7 and TET-inducible Ba/F3 cell lines

UT-7 cell lines were used for microarray experiments and western-blot analyses. The human hematopoietic (erythroid/megakaryoblastic) UT-7 parental cell line (UT-7/p) has been kindly provided to our lab by Dr Komatsu [12]. Its counterparts transduced with either native BCR-ABL1-p210 (UT-7/11 cells) or T315I-mutated-BCR-ABL1 (UT-7/T315I) resistant to first- and second-generation TKI and previously described and characterized by our group [13,14]. Doxycyclin-inducible BaF/p210 sin1.55 cell line has previously been described [15]. In this TET-OFF model, the addition of doxycycline in the cell culture medium turns off BCR-ABL1 expression.

### Patients and healthy donors

For *ENOX2* mRNA expression analysis, 36 CP-CML patients at diagnosis were tested (12 females and 24 males; median age 58.6 years, range 21.5-84.8). Median follow-up was 11.3 years (range 1.8-17.4). TKI discontinuation was initiated in 15 patients in deep molecular response after a median period of 5.7 years of treatment (S1 Table). Seven of them remained in DMR after a median of 6.4 years of treatment (range 5.3-10.4). A cohort of 27 healthy donors was used as controls. Concerning the quantification of ENOX2 protein in plasma samples, independent cohorts of 41 CML patients at diagnosis and 28 healthy controls were analyzed. This non-clinical study has been approved by the INSERM 935 Ethics committee (11 February 2014). All patients and healthy donors gave their informed consent in accordance with the Declaration of Helsinki.

### Transcriptome experiments

Transcriptome experiments were performed using UT-7/11 cells expressing BCR-ABL1 as compared to parental UT-7 cells. Total RNA (in triplicate) was extracted from UT-7 cells (expressing or not BCR-ABL1) after culture. RNA quantification was performed on Nanodrop (Thermo Fisher Scientific, Santa-Clara, CA), and sample quality was evaluated on the Bioanalyzer-2100 (Agilent Technologies, Santa Clara, CA). Transcriptome analysis was performed on the Affymetrix technologies platform (Affymetrix, Santa-Clara, CA). The results from triplicate samples of each experimental group were normalized with the RMA algorithm (Affymetrix).

### Quantitative RT-PCR assays

Total RNA from whole blood of CML patients at diagnosis and healthy donors was reverse transcribed using High Capacity cDNA Reverse Transcription Kit (Life Technologies, Foster City, CA), and qRT-PCR experiments were performed using 7900 Sequence Detection System (Life Technologies). TaqMan pre-developed assays reagent (Life Technologies) were used to quantify ENOX2 (Hs00197268_m1) and B2M (beta-2 microglobulin, Hs00187842_m1) mRNA transcripts. This housekeeping gene was used as an internal reference. Theoretically, Hs00197268_m1 TaqMan expression assay allows detection of all *ENOX2* mRNA splicing variants since the primers encompass exon 10 and 11 of the transcript variant 2 (NM_182314), taken as a reference (S2 and S3 Figs). Indeed, ENOX2 proteins could be the consequence of specific splicing variants promoting alternative downstream translation initiation sites [16]. Only *ENOX2* mRNA resulting from exon 7 skipping (also named exon 4 minus variant in another nomenclature) appears to be translated in a functional protein that localizes to the plasma membrane’s outer face of cancer cells. To quantify *ENOX2* mRNA, PCR reactions were prepared in duplicates in a final volume of 25 μL using the TaqMan Universal PCR Master Mix (Life Technologies). The %*ENOX2*/*B2M* was established using the ΔCt method.

### Western blotting

Cells were lysed in ice with RIPA buffer (NaCl 200mM, Tris pH=8 50mM, Nonidet P40 1%, deoxycholic acid 0.5%, SDS 0.05%, EDTA 2mM) supplemented with 100μM phenylmethylsulfonyl fluoride, 1mM sodium fluoride, 1mM sodium orthovanadate (Sigma-Aldrich, St. Louis, MO). Protein separation was performed by electrophoresis on 8% polyacrylamide gel under denaturing conditions. Proteins were then transferred on PVDF membrane pre-activated in methanol. After saturation with TBS-Tween-BSA for 1 hour, membranes were hybridized with primary and HRP coupled secondary antibodies (New England Biolabs, Ipswich, MA). Membranes were revealed by chemiluminescence with SuperSignal West Dura or Femto reagents and data acquired using the G-BOX-iChemi Chemiluminescence Image Capture system (Syngene, Frederick, MD). The ENOX2 antibody used for these experiments was purchased from LSBio (LS – C346209, LifeSpan BioSciences Inc. Biotechnology. Seattle, WA). The ABL1 antibody, for the detection of translocated and non-translocated ABL1 protein, was obtained from Santa Cruz Biotechnology (Santa Cruz, CA), and beta-ACTIN (Sigma-Aldrich) was run as a control. All antibodies were used at the dilution recommended by the manufacturer.

### ENOX2 protein concentration in blood plasma

ENOX2 proteins can be shed into extracellular fluids, in which they can be detected [17]. In this study, ENOX2 protein quantitation in blood plasma was performed using the human ecto-NOX disulfide-thiol exchanger 2 ELISA Kit (MBS 943476, MyBioSource Inc, San Diego, CA) according to the manufacturer’s guidelines.

### Transcriptome dataset GSE4170

Using DNA microarray, Radich et al aimed to compare gene expression between chronic (<10% blasts), accelerated (10–30% blasts), and blast phase (>30% blasts) by analyzing 91 cases of CML using normal immature CD34+ cells as a reference [18]. Rosetta/Merck Human 25k v2.2.1 data matrix of normalized log-ratios from GSE4170 was downloaded on the Gene Expression Omnibus (GEO) website and annotated with the annotation platform GPL2029 to be reanalyzed, focusing on *ENOX2* mRNA expression.

### Bioinformatics microarray data analysis

Differentially expressed genes between the three phases of CML (GEO dataset GSE4170) were determined by Pavlidis template matching algorithm with a positive R correlation coefficient greater than 0.80 and p-value threshold less than 1E-6 [19]. Heatmap and parallel coordinate plot were carried out with the made4 R package [20] and the GGally R-package. The expression profile correlated to *ENOX2* in CML cells was used to generate functional enrichment with Go-Elite Standalone software version 1.2 on the Gene Ontology Biological Process database included in Homo sapiens EnsMart77Plus (Ensembl – Biomart) update [21]. Unsupervised principal component analysis was performed on correlated gene expression profile with R package FactoMiner and p-value was calculated by group discrimination on the first principal component axis. The functional interaction network was built with functional relations identified during enrichment analysis with Cytoscape software version 3.2.1 [22].

### Scatter dot plots, boxplots, and statistical analysis

Data from ENOX2 gene expression or protein concentration in plasma were expressed as scatter dot plots or boxplots with medians. They were generated using GraphPad Prism version 8.0 (GraphPad Software, San Diego, CA). Welch’s t-test determined statistical significance between data groups. Differences are considered significant at p<0.05.

## Results

### *ENOX2* mRNA is overexpressed in BCR-ABL1-expressing UT-7 cell lines

We conducted a transcriptome assay to identify genes up- or down-regulated in *BCR-ABL1*-expressing UT-7 cell lines. To this end, we compared the UT-7/11 cell line (n=3), which expressed high levels of BCR-ABL1 protein, to parental UT-7 (UT7/p, n=3). We focused on ENOX2 mRNA, which was not previously known to be over-expressed in BCR-ABL1-positive leukemia. In our experiments, *ENOX2* was significantly up-regulated (X 2.4, p=0.0012) in the *BCR-ABL1*-expressing cell line (Fig 1).

**Fig 1.**
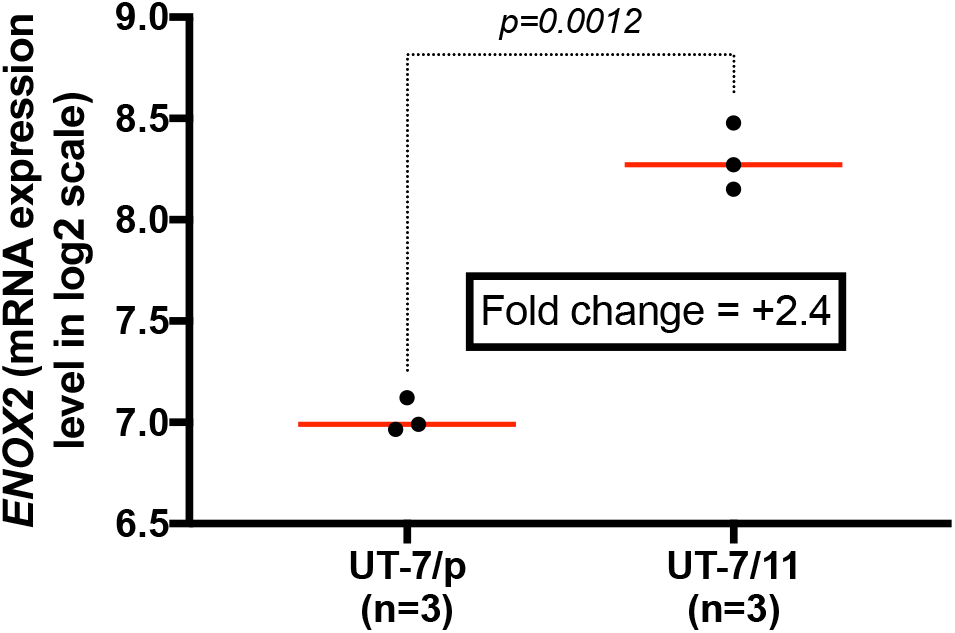
*ENOX2* up-regulation in BCR-ABL1-positive UT-7/11 cell line. Significant overexpression of *ENOX2* mRNA is observed in UT-7/11 (transfected by *BCR-ABL1*) compared with UT-7 controls. P-values were calculated with a two-sided Welch’s t-test.

### *ENOX2* mRNA expression is increased in primary cells from patients at diagnosis

To validate these preliminary results obtained in UT7 cells transduced with *BCR-ABL1*, we examined *ENOX2* mRNA expression by qRT-PCR in the blood samples from a cohort of CP-CML patients at diagnosis (n=36) and a series of healthy donors (n=27). As shown in Fig 2, levels of *ENOX2* mRNA were significantly increased in samples from CML patients at diagnosis (p<0.0001) compared to healthy donors, with a fold change of 4.75. In this cohort, we did not find any correlation between *ENOX2* mRNA expression and Sokal score, or patient outcomes (achievement of a deep molecular response or sustained TFR after TKI withdrawal).

**Fig 2.**
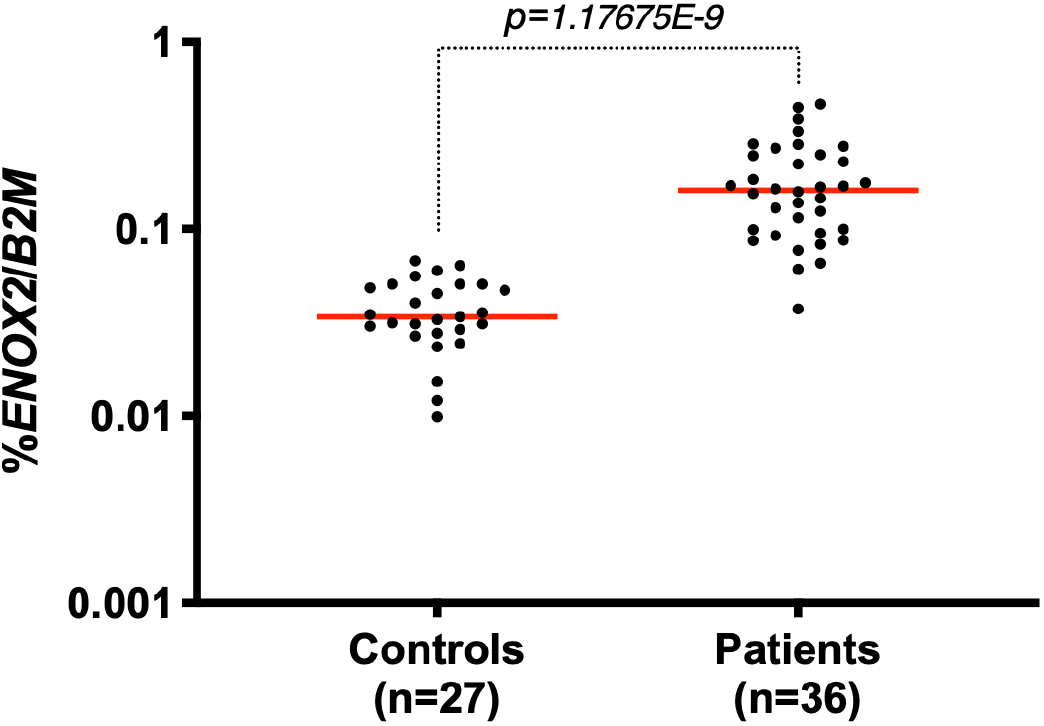
*ENOX2* mRNA is overexpressed in primary cells from CML patients. The scatter dot plot shows *ENOX2* mRNA expression in whole blood samples from CML patients at diagnosis (n = 36) compared with healthy donors (n=27). P-values were calculated with a two-sided Welch’s t-test.

### BCR-ABL1 induces the production of ENOX2 protein in UT7 and TET-inducible Ba/F3 BCR-ABL1 cell lines

Three UT-7 cell lines were first used to analyze the presence of ENOX2 protein by western-blots (parental UT-7, UT-7/11 and UT-7/T315I). We generated UT-7/11 and UT-7/T315I cell lines several years ago by retroviral transduction of wild-type or T315I-mutated *BCR-ABL1* in the GM-CSF-dependent parental UT-7 cell line. In these two cell line models, the *BCR-ABL1* expression renders these cells GM-CSF-independent [13]. As can be expected, Fig 3a confirms the presence of BCR-ABL1 protein in UT-7/11 and UT-7/T315I cell lines, in contrast to parental UT-7. Concerning ENOX2, high protein levels were produced in the *BCR-ABL1*-expressing UT-7 cell line (native UT-7/11 and mutated-UT-7/T315I) compared to UT-7/p control (Fig 3b).

**Fig 3.**
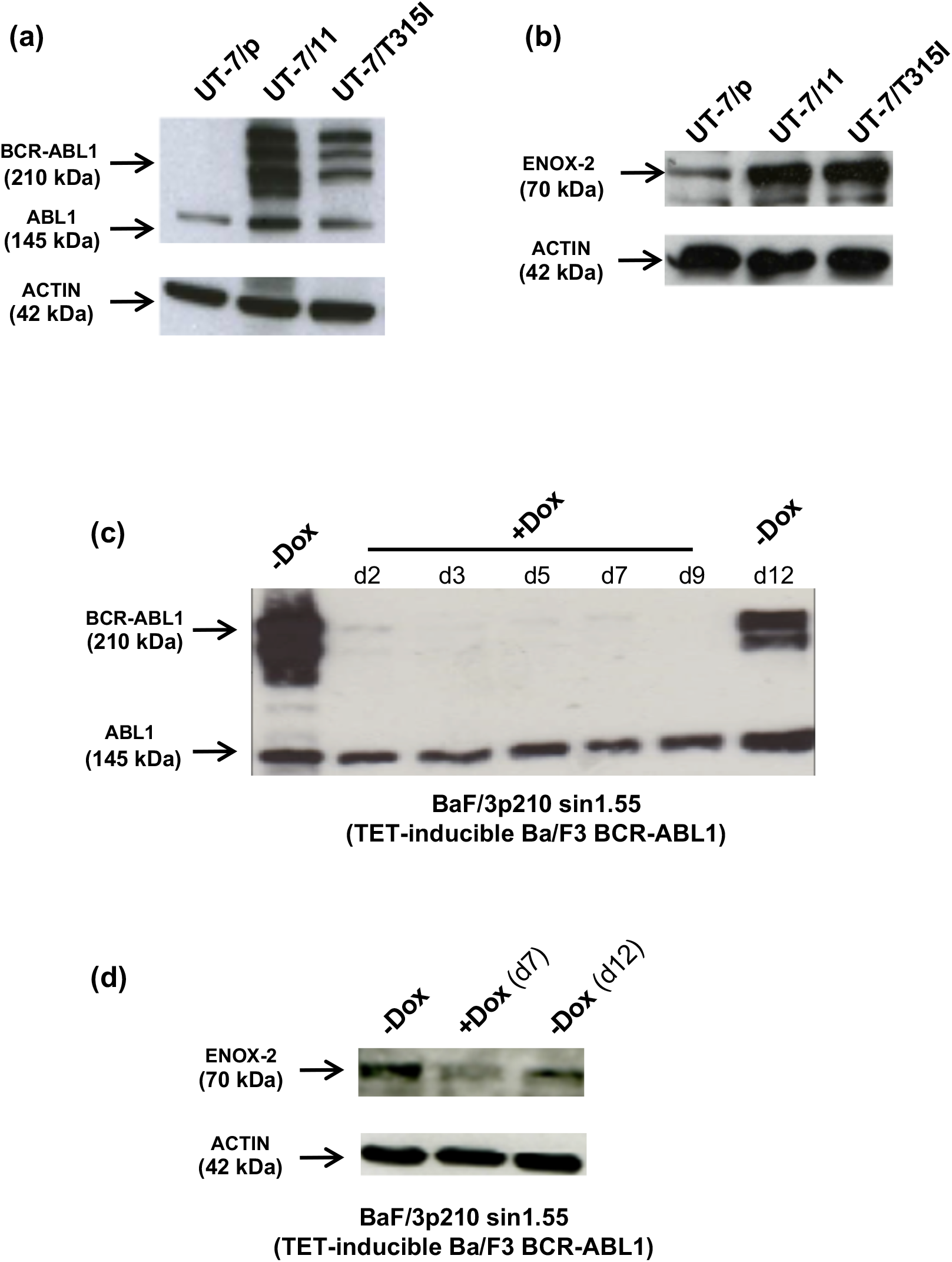
ENOX2 protein is up-regulated by BCR-ABL1 in UT-7/11 and TET-inducible Ba/F3 cell lines. (**a**) Western blot control of BCR-ABL1 induction after the transfection of the *BCR-ABL1* fusion oncogene in the UT-7/11 cell line (non-mutated UT-7/11 and mutated UT7/T315I) as compared to parental UT-7. (**b**) High ENOX2 up-regulation is observed by western blot analysis in transfected BCR-ABL1 UT-7 cells (non-mutated UT-7/11 and mutated UT7/T315I). **(c**) Western blot verification of the inducible Ba/F3 (BaF/p210 sin1.55) cell line expressing BCR-ABL1 under the control of a TET promoter. As shown, cells were cultured in the presence of doxycycline for 2, 3, 5, 7, and 9 days. As can be seen in the Figure, upon washing out doxycycline from the cell medium, BCR-ABL1 is re-expressed at day 12. (**d**) ENOX2 up-regulation in the inducible Ba/F3 cell line is dependent on the presence of BCR-ABL1. As observed in the Figure, upon washing out doxycycline from the cell medium, ENOX2 is re-expressed at day 12.

To ensure that ENOX2 overexpression would be related to the presence of BCR-ABL1, we used a Ba/F3 cell line transduced with *BCR-ABL1* under the control of the TET promoter (BaF/p210 sin1.55). This inducible model was appropriate insofar as the BCR-ABL1 protein was inhibited by doxycycline (Fig 3c). Decreased ENOX2 protein expression was also observed in response to doxycycline added to the culture medium (Fig 3d). As can be seen in Figs 3c and 3d, BCR-ABL1 and ENOX2 are re-expressed at day 12 upon washing out doxycycline from the cell medium. Consequently, ENOX2 overexpression appeared to be related to the presence of BCR-ABL1 insofar as inhibition of BCR-ABL1 expression in the TET-inducible Ba/F3 model led to a reduction in ENOX2 protein synthesis.

Finally, we wondered whether the expression of ENOX2 in CML cells was a tyrosine kinase-dependent event. We have previously demonstrated the reduction of phospho-Tyr BCR-ABL1 protein upon Imatinib treatment in *BCR-ABL1*-expressing UT7 cells [23]. In this study, western blot experiments showed that ENOX2 protein expression was reduced in the UT7/11 cell line treated with Imatinib 1 microM for 6, 18, and 24 hours (Fig 4a). Same experiments performed on UT-7/p cell line showed no modification of ENOX2 expression (Fig 4b).

**Fig 4.**
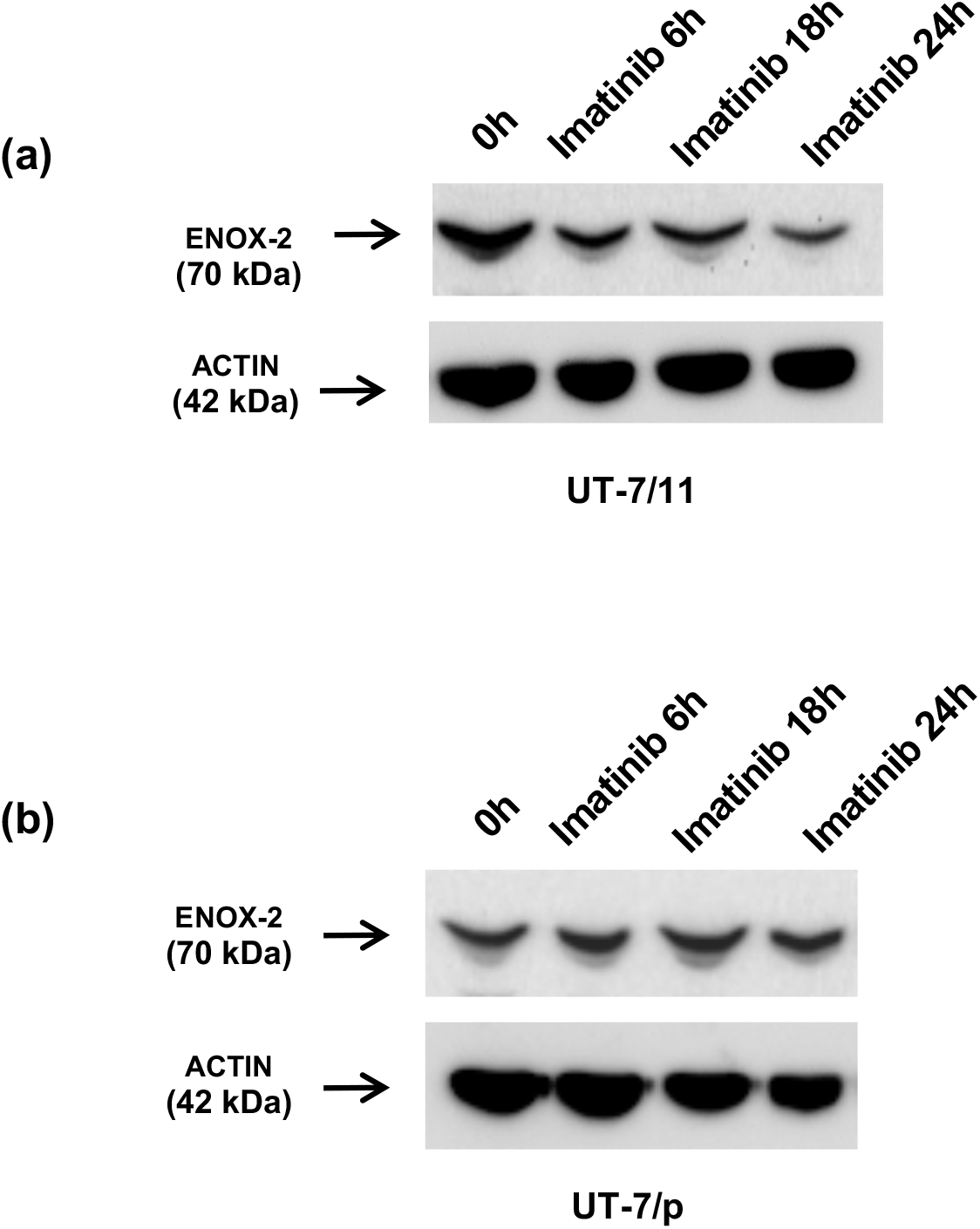
ENOX2 protein up-regulation depends on the BCR-ABL1 tyrosine kinase-activity. **a**) Western blot analysis performed in the UT-7/11 cell line under tyrosine kinase inhibition conditions (Imatinib 1 microM for 6, 18, 24 hours) shows the reduction of the ENOX2 protein expression, suggesting that its expression is dependent on the kinase activity of BCR-ABL1. **b**) Western blot analysis performed in the UT-7/p cell line as control showing no modification of ENOX2 expression upon inhibition of BCR-ABL1 tyrosine kinase activity.

### ENOX2 protein levels are significantly increased in the plasma of CML patients

As ENOX2 protein can be released from tumor cells into extracellular fluids, we determined its concentration in the plasma of CML patients at diagnosis (n=41) compared to healthy controls (n=28) using an ELISA method. As presented in Fig 5, a significant increase in plasma ENOX2 protein levels (p<0.0001) was shown in CP-CML patients at diagnosis before TKI therapy. The extended frequency distribution of ENOX2 protein levels is undoubtedly due to heterogeneity between patients. Furthermore, no correlation was found between ENOX2 protein levels in the plasma and white blood cell (WBC) count at diagnosis (S4 Fig).

**Fig 5.**
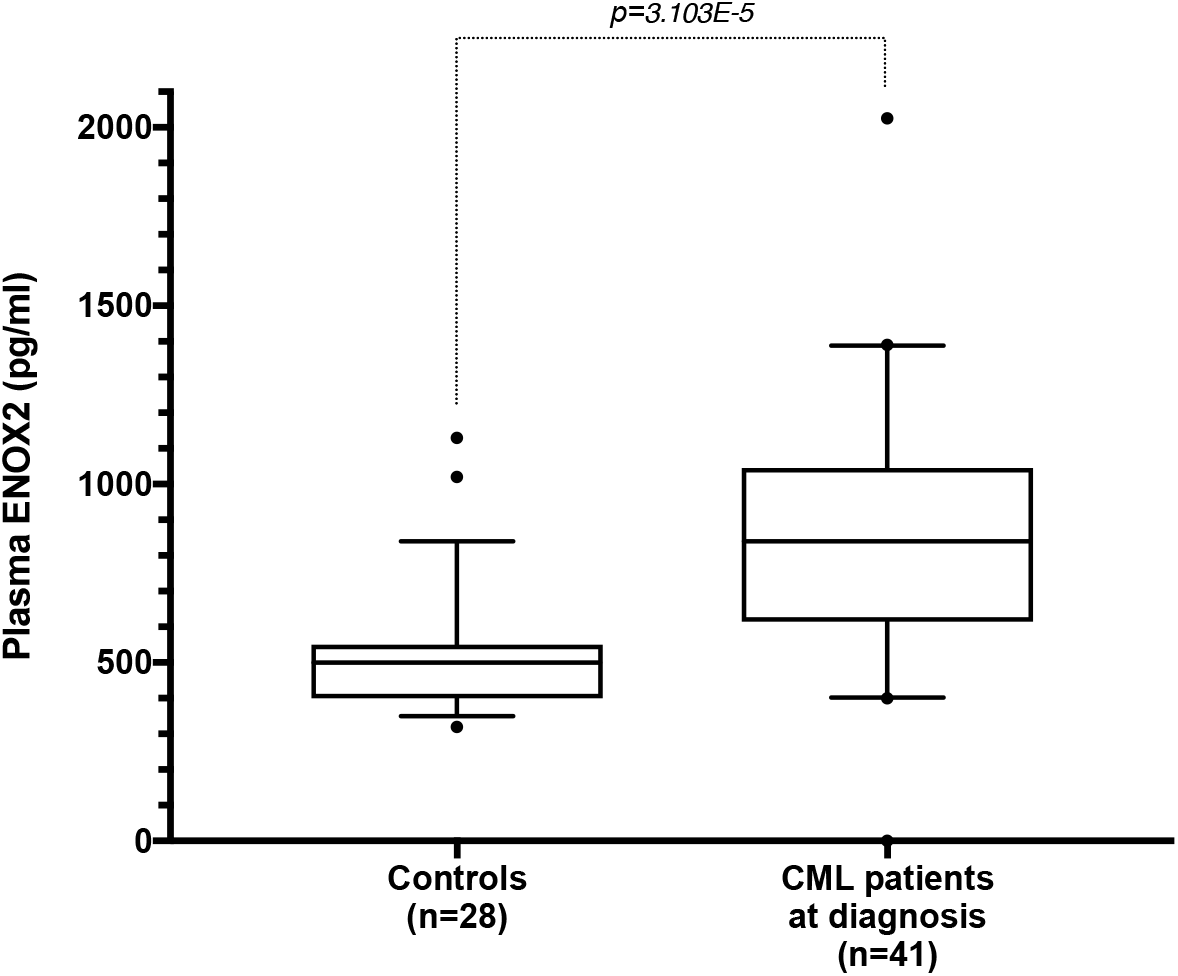
Secreted ENOX2 protein is detected in blood from CP-CML patients at diagnosis. The boxplot shows a high ENOX2 protein level in blood plasma from CP-CML patients at diagnosis (n = 41) compared with healthy donors (n=28). The median and the 5%, 25%, 75%, and 95% percentiles are shown in boxplot diagrams. Dots correspond to observations outside the range of adjacent values. P-values were calculated with a two-sided Welch’s t-test.

### *ENOX2* mRNA overexpression is characteristic of the chronic phase of CML

We reanalyzed a publicly available transcriptome dataset from CML patients during the different phases of their disease (GSE4170). In this work, the authors performed microarray analyses to evaluate CML patients’ gene expression profiles in the chronic, accelerated, and blast phases. *ENOX2* mRNA expression data were retrieved from the GE0 database and analyzed independently. For each patient, *ENOX2* mRNA expression was shown using normal immature CD34^+^ cells as reference (median *ENOX2* expression of normal CD34+ cells was subtracted from each sample). As shown in Fig 6, *ENOX2* mRNA up-regulation was observed in this independent study for all CP-CML patients. In addition, *ENOX2* was significantly overexpressed in CP-CML patients, as compared to patients in the accelerated (p<0.0001) and blast phases (p<0.0001). Interestingly, *ENOX2* expression in the accelerated and blast crises is very similar to that of normal CD34+ cells and appears heterogeneous.

**Fig 6.**
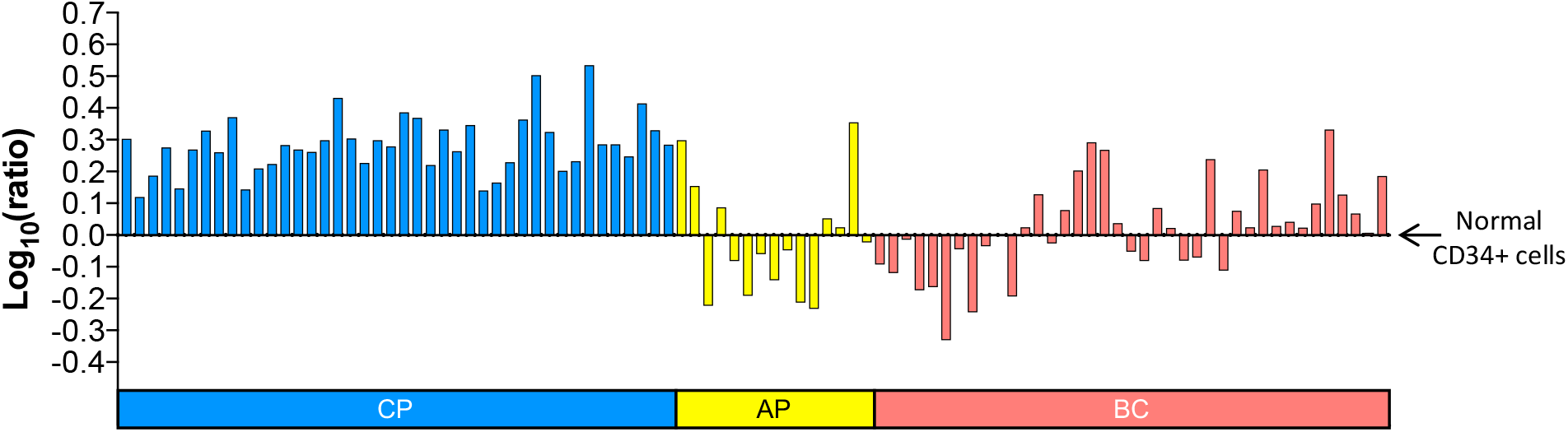
Relative *ENOX2* gene expression during the different phases of CML using the GSE4170 GEO dataset. *ENOX2* mRNA relative expressions in CML leukemic cells corresponding to patients in chronic phase (CP), accelerated phase (AP), and blast crisis (BC) were compared to normal immature CD34^+^ cells. P-values were calculated with a two-sided Welch’s t-test.

### The mRNA expression of *ENOX2* and related genes distinguishes the chronic phase of CML from advanced phases

Exploiting the GSE4170 transcriptome dataset, we then tried to discover genes positively correlated to *ENOX2* mRNA expression and to highlight a potential link between the *ENOX2* transcriptional program and BCR-ABL1-mediated leukemogenesis. Pavlidis template matching algorithm used with *ENOX2* as a predictor highlighted 301 related genes with a positive r correlation coefficient greater than 0.80 and a p-value threshold less than p<1E-6 (S2 Table). The high level of correlation to *ENOX2* mRNA expression is illustrated for eight potentially relevant protein-coding genes (*EPHB3*, *HEYL*, *ERK1*, *PlGF*, *FAK*, *RHOG*, *THY1,* and *TRIB1*) in S5 Fig. Heatmap and parallel coordinate plot revealed that the high mRNA expression level of these 301 genes was characteristic of the chronic phase of CML (Fig 7a). Unsupervised principal component analysis performed with *ENOX2* pattern matching gene expression profile allowed highly significant discrimination of the chronic phase from accelerated and blast phases (Fig 7b, p<0.0001).

**Fig 7.**
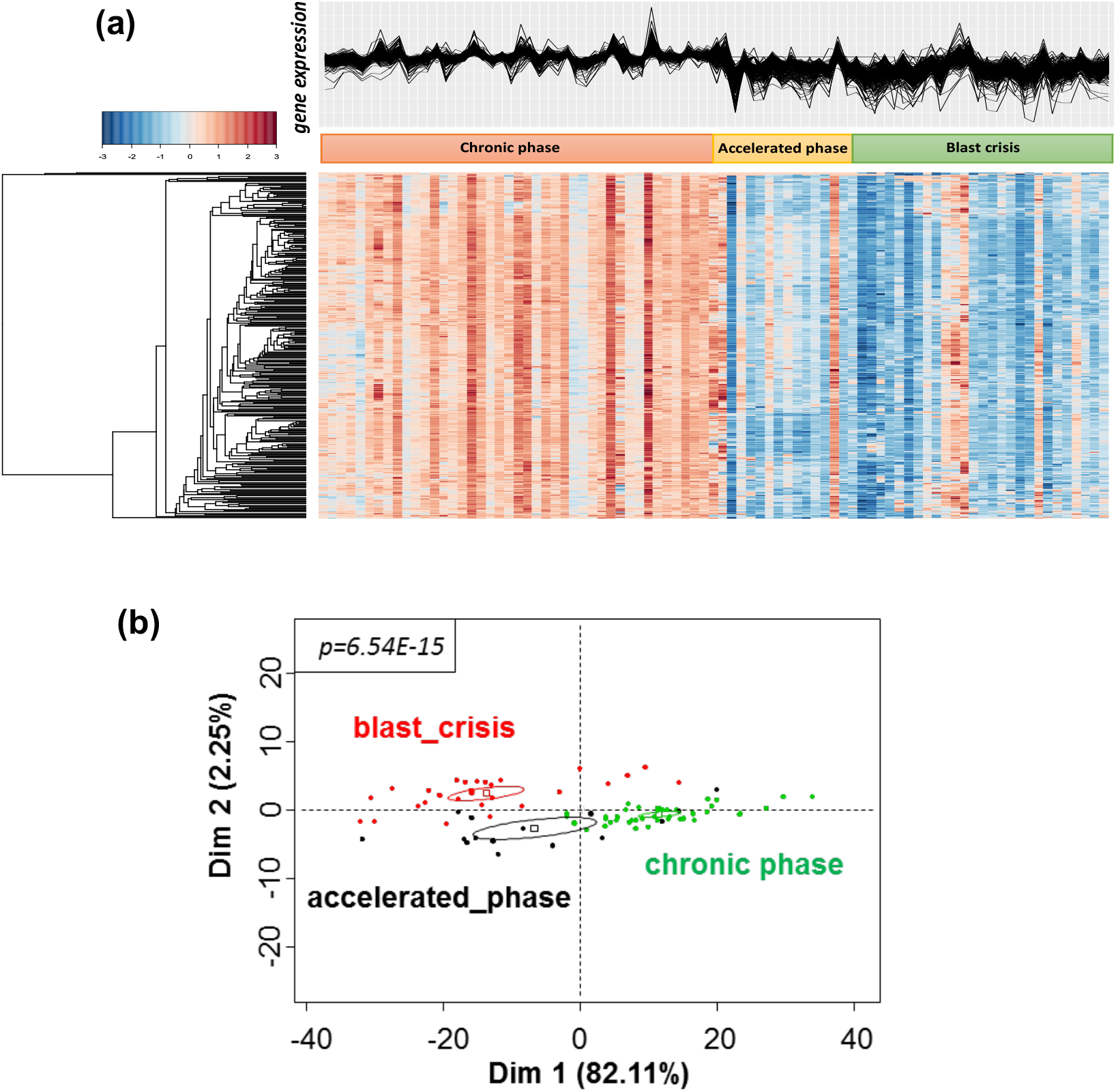
The mRNA expression of *ENOX2* and related genes allows discrimination of the chronic phase of CML from the more advanced phases of the disease. (**a**) Heatmap and parallel coordinate plot revealed 301 genes positively correlated to *ENOX2* mRNA expression. Their high mRNA expression level appears to be characteristic of the chronic phase of CML. (**b**) Unsupervised principal component analysis performed with ENOX2 pattern matching gene expression profile distinguishes the chronic phase of CML from the accelerated and blast phases. Correlation p-value was evaluated for phase discrimination on the first principal axis; ellipses around the barycenter were estimated with 75% confidence.

### Several proteins of a potential ENOX2 network can be involved in the CML context in crucial biological processes

Functional enrichment performed with ENOX2 pattern matching on the Gene Ontology Biological Process database emphasized 49 genes (out of 301) known to be involved in essential cell processes (Fig 8a). In the context of *ENOX2* mRNA overexpression in CP-CML, several major biological functions appear to be activated: angiogenesis, NOTCH signaling, cell morphogenesis differentiation, circadian rhythm, RAS signaling, cell proliferation, G-protein receptor pathways, integrin-mediated signaling, carbohydrate homeostasis, stress-activated protein kinase, and RHO GTPase activities (Fig 8b, S3 Table).

**Fig 8.**
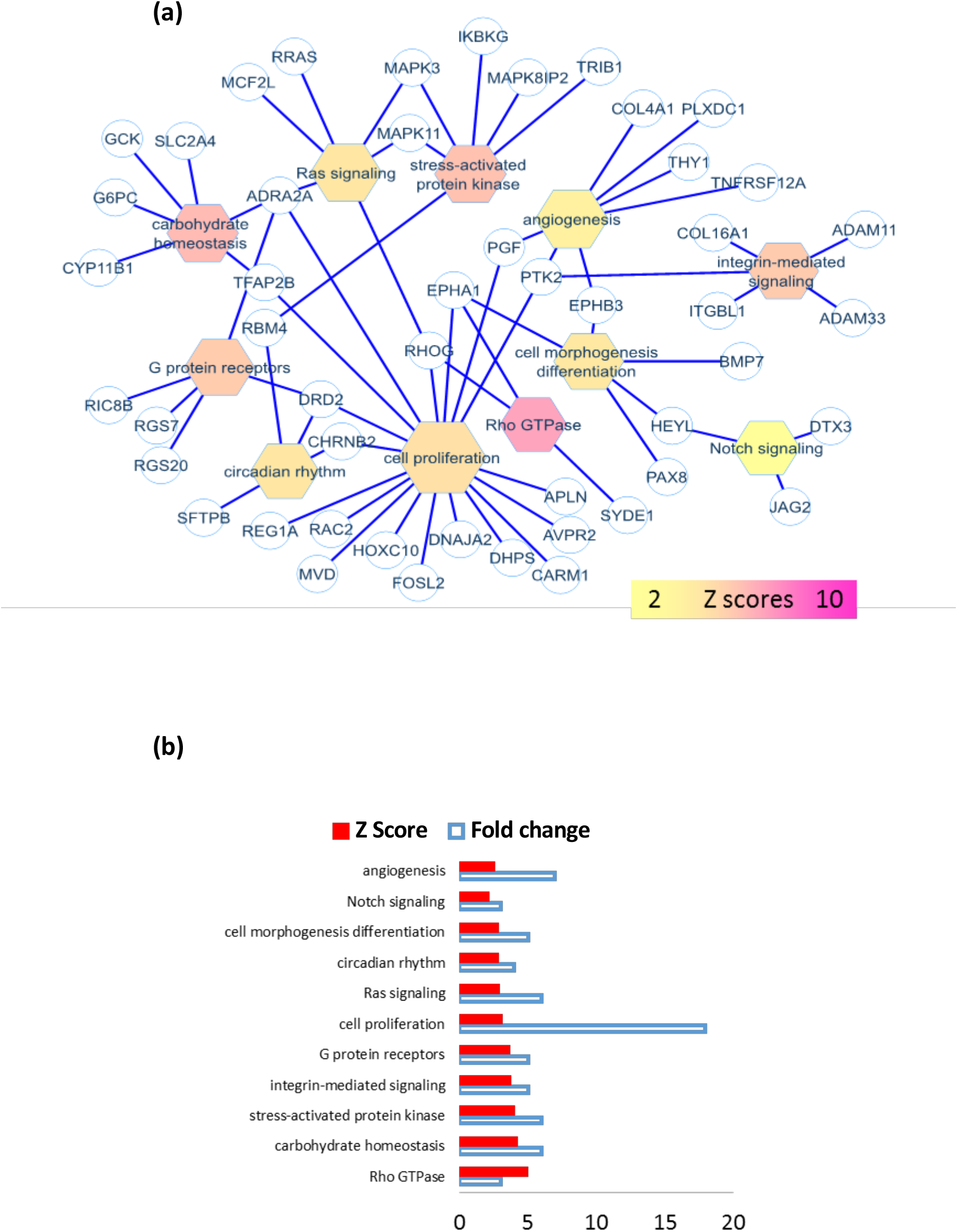
A potential network correlated to *ENOX2* gene expression is highlighted in CML cells. (**a**) Functional enrichment performed with *ENOX2* pattern matching on the Gene Ontology Biological Process database reveals 49 genes (out of 301) known to be involved in essential cellular processes. Results are presented as a Cytoscape network view showing both biological functions and related genes. The color of nodes in the network is relative to the Z-scores obtained during functional enrichment (from yellow for low Z-scores to purple for high Z-scores), the size of function nodes is relative to the number of genes enriched in the corresponding function mapped on the network. (**b**) Bar plots represent both the Z scores and the gene expression fold changes for the different molecular functions defined by the Gene Ontology Biological Process database.

## Discussion

The involvement of ENOX2 in BCR-ABL1-positive leukemias, as indicated by PubMed searches, has yet to be shown. In this work, we established that *ENOX2* mRNA was overexpressed in both experimental models of BCR-ABL1-induced cell transformation and primary leukemic cells from newly diagnosed CP-CML patients. Regarding the *in vitro* model, we did not use CML cell lines derived from patients in blast crisis, such as the erythrocytic K562, the myelocytic KCL-22 or the erythro-megakaryocytic LAMA-84, since they present an extreme variability in their characteristics of cell clones and in their BCR-ABL1 expression. Moreover, there is no counterpart of these cells without *BCR-ABL1* expression (as opposed to UT7 model). Western-blots analyses confirmed the presence of higher levels of ENOX2 protein in *BCR-ABL1*-expressing murine and human cell lines. This overexpression is directly influenced by the presence of BCR-ABL1 oncoprotein and related to its constitutive tyrosine kinase activity. Using a large number of plasma samples from both CML patients and healthy donors, we went on to show a significant increase in plasma ENOX2 levels in CML patients at diagnosis. Reanalyzing a transcriptomic dataset from a previously reported gene profiling study [18], we observed that *ENOX2* mRNA expression is restricted to the chronic phase of the disease. Nevertheless, we could not determine here whether ENOX2 is a direct or an indirect target of BCR-ABL1.

Several lines of experimental data indicate that ENOX2 protein expression in adult cells appears to be restricted to cancer cells. Although *ENOX2* mRNA was detectable in both non-malignant and malignant cells, the *ENOX2* gene would be translated only during early embryogenesis and cancer development. Therefore, ENOX2 physiologic function appears to be restricted to the embryonic period [24], in which this oncofetal protein located at the plasma membrane has an oscillating enzymatic activity. Little is known about the reappearance and the oncogenic function of ENOX2 in adult cells. However, it has been shown that ENOX2 proteins are constitutively activated in cancers and could promote cell proliferation [25–27].

As they are not firmly anchored into the cell membrane, ENOX2 proteins can be released into extracellular fluids. This circulating form has been detected in the sera of patients suffering from various tumors [28], whereas it is present only at very low levels in healthy subjects [29]. A two-dimensional gel electrophoretic separation (ONCOblot) reveals unique profiles that are seemingly specific to a type of cancer [17,28]. In the present study, high levels of circulating ENOX2 protein detected in plasma from CML patients at diagnosis are consistent with these data.

*In silico* reanalyses suggest that mRNA expression of several genes is positively correlated to *ENOX2* expression in a BCR-ABL1 context, thereby highlighting critical biological functions (angiogenesis, NOTCH signaling, cell morphogenesis differentiation, circadian rhythm, RAS signaling, cell proliferation, G-protein receptor pathways, integrin-mediated signaling, carbohydrate homeostasis, stress-activated protein kinase, and RHO GTPase activities). These data are based on mRNA expression and not on the presence/activity of the proteins. However, these results on a substantial number of individual genes are highly significant, and it is quite unlikely that all these genes are not translated into functional proteins. Consequently, high *ENOX2* mRNA expression probably goes along with the activation of cell proliferation and differentiation through genes encoding proteins from different pathways. ENOX2 overexpression has been linked to cell proliferation, migration, and increased expression of mesenchymal markers in some cancers [27]. Interestingly, these data are in line with the fundamental characteristics of CML dysregulation [30].

Among the genes found to be positively correlated with *ENOX2* mRNA expression in a CML context, some deserve further discussion. RAC2 GTPases are known to be critical regulators of BCR-ABL1-mediated leukemogenesis [31]. Activation of the MAP-kinase ERK through the RAS-induced pathway has been implicated in cell proliferation and differentiation in K562 cells [32]. The focal adhesion kinase (FAK), phosphorylated by BCR-ABL1, plays a crucial role in CML cells insofar as its silencing inhibits the leukemogenesis process [33]. The role of TRIB1 (tribbles homolog 1) serine/threonine kinase-like protein in CML pathogenesis has also been pointed out [34]. On the other hand, NOTCH1 has been shown to interact with BCR-ABL1 in CD34+ cells from CP-CML patients [35]. Finally, *BCR-ABL1*-expressing cells could secrete angiogenic factors [36]. In this respect, it has been shown that the BCR-ABL1 oncoprotein induces the production of placental growth factor (PGF or PlGF) by bone marrow stromal cells to support *in situ* angiogenesis and promote cell proliferation [37].

All these data suggest that ENOX2 could play a role in CML’s pathogenesis in a BCR-ABL1-dependent manner. In addition to this potential biological feature, could ENOX2 be a surrogate marker or even a therapeutic target? Based on our experiments, we can say that ENOX2 is the first secreted biomarker described in CML. On the other hand, during our study we did not find any relationship between *ENOX2* mRNA expression or ENOX2 plasma levels and the clinical course or the most relevant biological parameters. Some anti-oxidant agents (capsaicin, omega-3 polyunsaturated fatty acids, or synthetic isoflavone) could exert an anti-tumor effect by inhibiting ENOX2 enzymatic activity [38–40]. All of these agents (and perhaps others) could represent a potential ENOX2-targeted therapy in malignant diseases. However, in CML, a majority of patients treated with TKIs achieve a sustained molecular response. Therefore, in most cases, CML treatment does not require additional drugs. Nevertheless, two circumstances may require other therapeutic options: complete resistance to all available TKIs, or the persistence of quiescent leukemic stem cells [41]. The potential interest of ENOX2 as a druggable target in these contexts calls for further exploration.

## Conclusions

It is now well-established that ENOX2 protein is present during the embryonic period, almost absent in normal adult cells, and reappears in most cancer cells. In physiological conditions, ENOX2 fulfills functions essential to the development of early embryos. During pathological circumstances, the same enzymatic functions could contribute to growth deregulation of cancer cells. Based on our results, we propose that BCR-ABL1 up-regulates ENOX2 in the chronic phase of CML. To the best of our knowledge, the association between the reactivation of ENOX2 in adult cells and deregulated tyrosine kinase activity was never previously observed. That said, a link between BCR-ABL1 and the tumor-associated NADH oxidase ENOX2 has not been established. The hypothesis that some downstream BCR-ABL1 substrates could activate an irrelevant translation mechanism of ENOX2 variant transcripts can be put forward.

## Acknowledgements

The English of the manuscript was reviewed by Jeffrey Arsham, an American medical translator.

## Author’s contributions

Conceptualization, A.B.G., J.C.C. and A.G.T.; methodology, S.B.K., M.V. and N.S.; experiments, S.B.K., M.V. and N.S.; bioinformatic analysis, C.D.; validation, J.C.C., A.B.G. and A.G.T.; clinical investigation, E.C., H.G.A, H.G.B and A.G.T.; writing—original draft preparation, J.C.C. and A.G.T.; writing—review and editing, J.C.C., N.S., E.C., A.B.G and A.G.T.; supervision, A.G.T. All authors have read and agreed to the published version of the manuscript.

## Supporting information

**S1 Fig:**
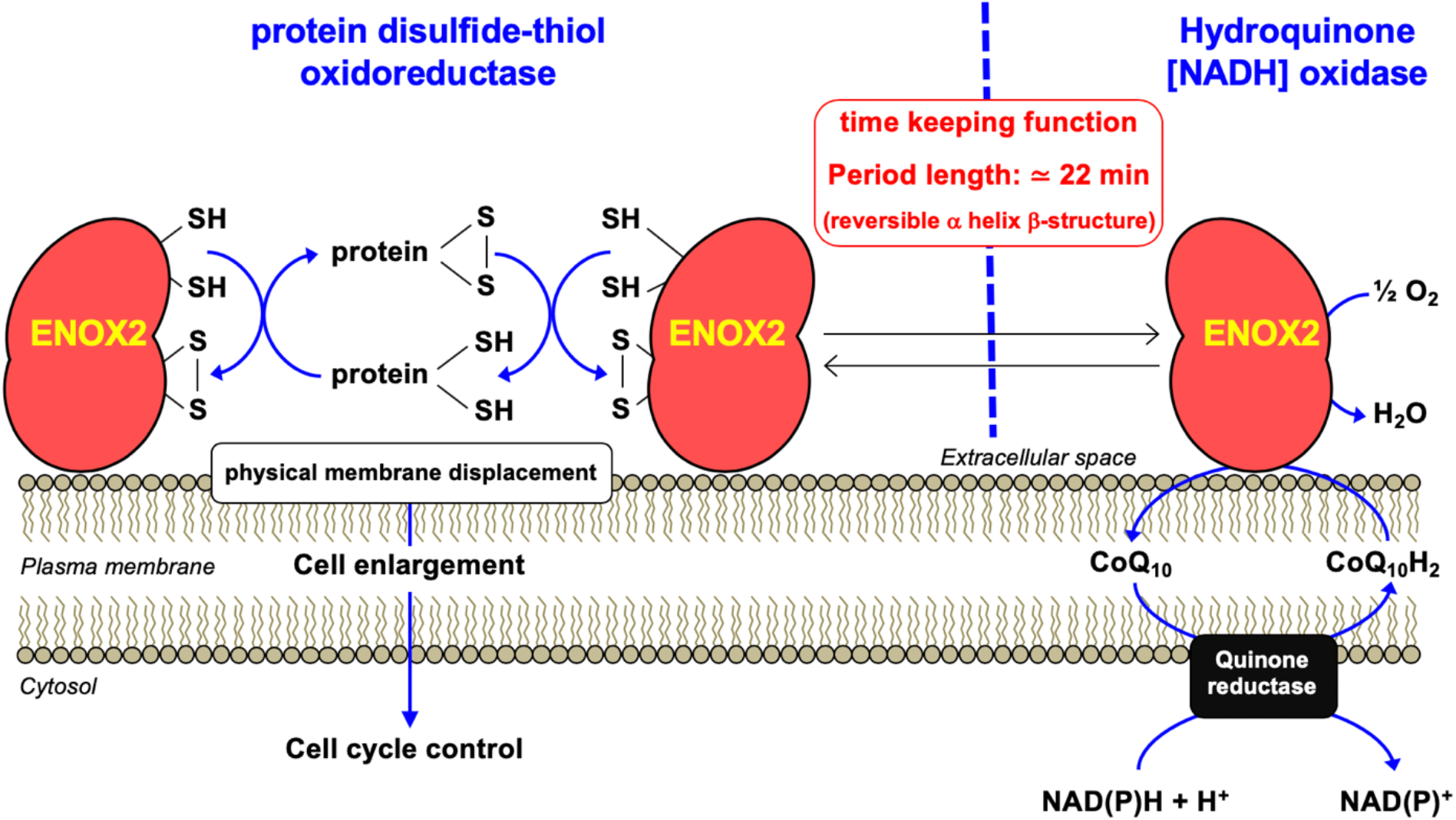
Schematic representation of ENOX2 dimer functions at the plasma membrane of a cancer cell. ENOX2 combines two main enzymatic oscillatory activities (hydroquinone NADH oxidase and protein disulfide-thiol oxidoreductase) that alternate with a period length of approximately 22 min, generating an ultradian cellular biological clock of 22 hours. According to Morré DJ, Morré DM. ECTO-NOX Proteins, Growth, Cancer, and Aging. Springer (2013).

**S2 Fig:**
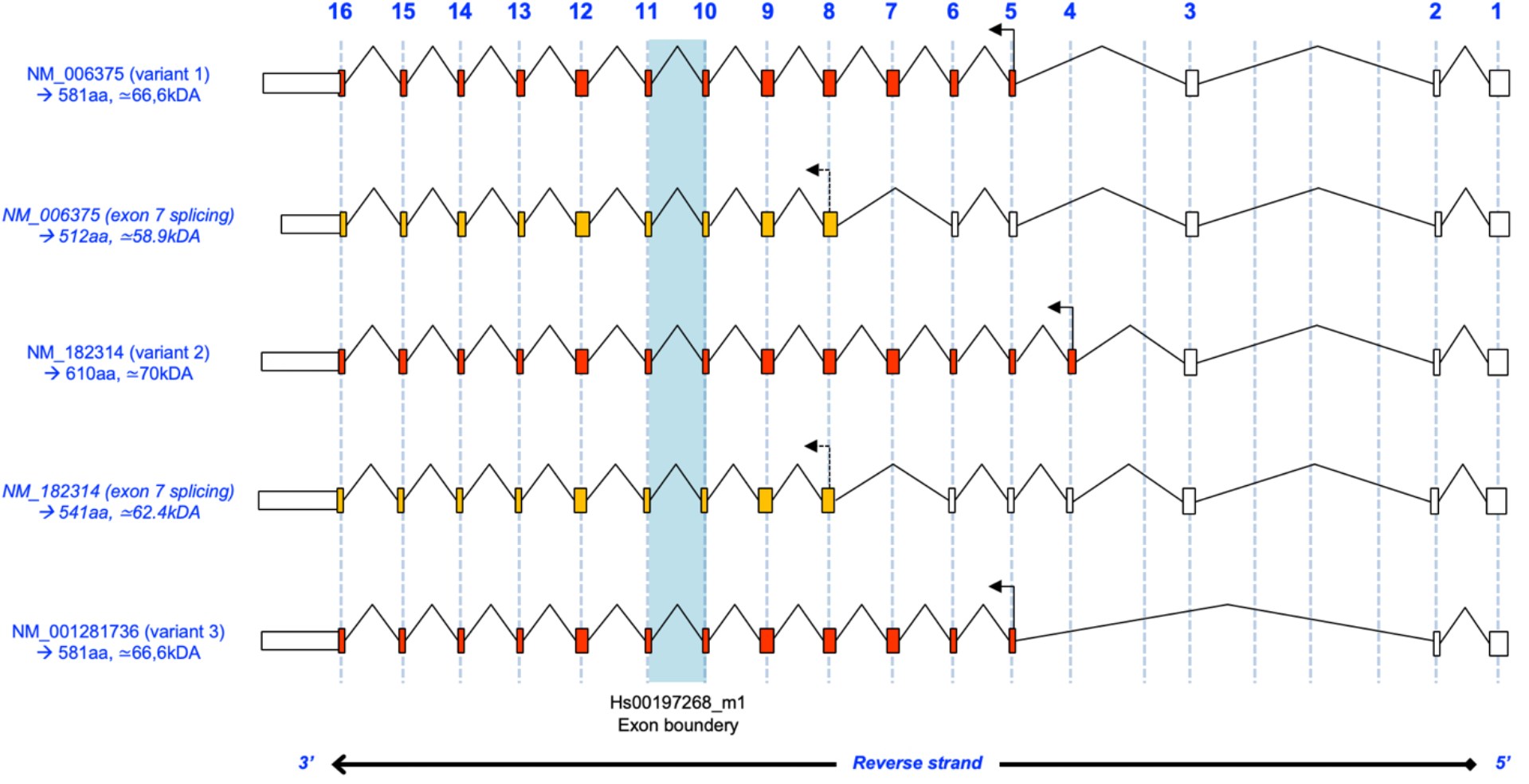
Schematic representation of some mature transcript types (as examples) and alternative splicing observed in the *ENOX2* gene. Data came from REFSEQ mRNAs (NM_182314 taken as reference for exon numbering). The blue bar represents exon 10-11 boundaries for qRT-PCR experiments using *ENOX2* TaqMan pre-developed assay reagent (Hs00197268_m1). Theoretical derived molecular weights of proteins translated from the open reading frame of the different mRNA isoforms are given as an indication. Exons in red and orange correspond to full-length cDNA and exons potentially translated in case of exon 7 skipping (alternative translation initiation sites), respectively.

**S3 Fig:**
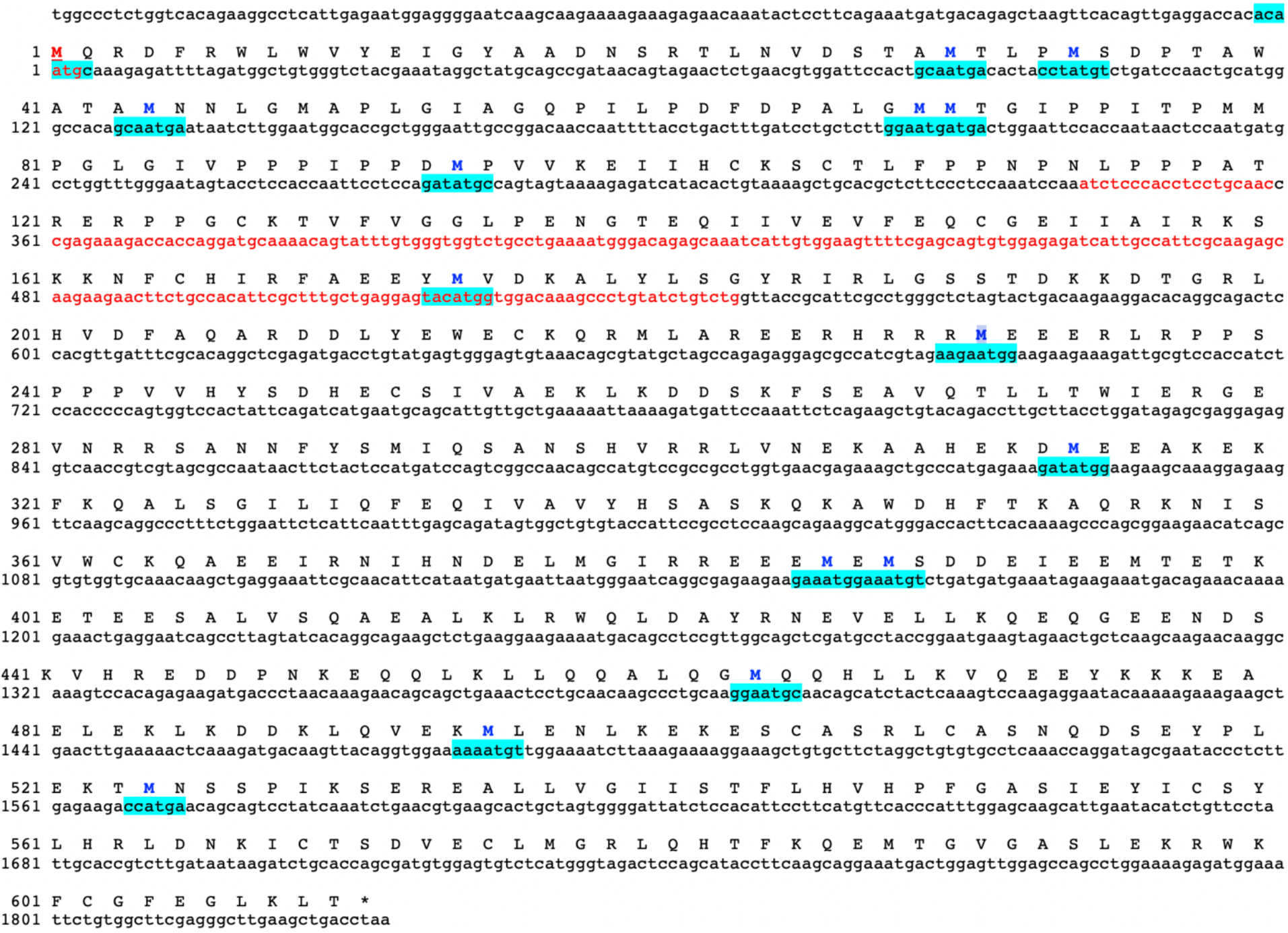
Amino-acid and nucleotide sequences of full-length *ENOX2*. Amino-acid and nucleotide numeration correspond to the theoretical protein of 610aa (70kDa). The sequences underlined in blue indicate the most likely Kozak consensus for protein translation initiation. Sequences in red correspond to exon 7 (exon 4 in another nomenclature). In the case of exon 7 skipping (NM_183314 numbering), alternative translation initiation sites, such as the M^231^, initiate the production of various tumor-associated ENOX2 proteins of lower molecular weight.

**S4 Fig:**
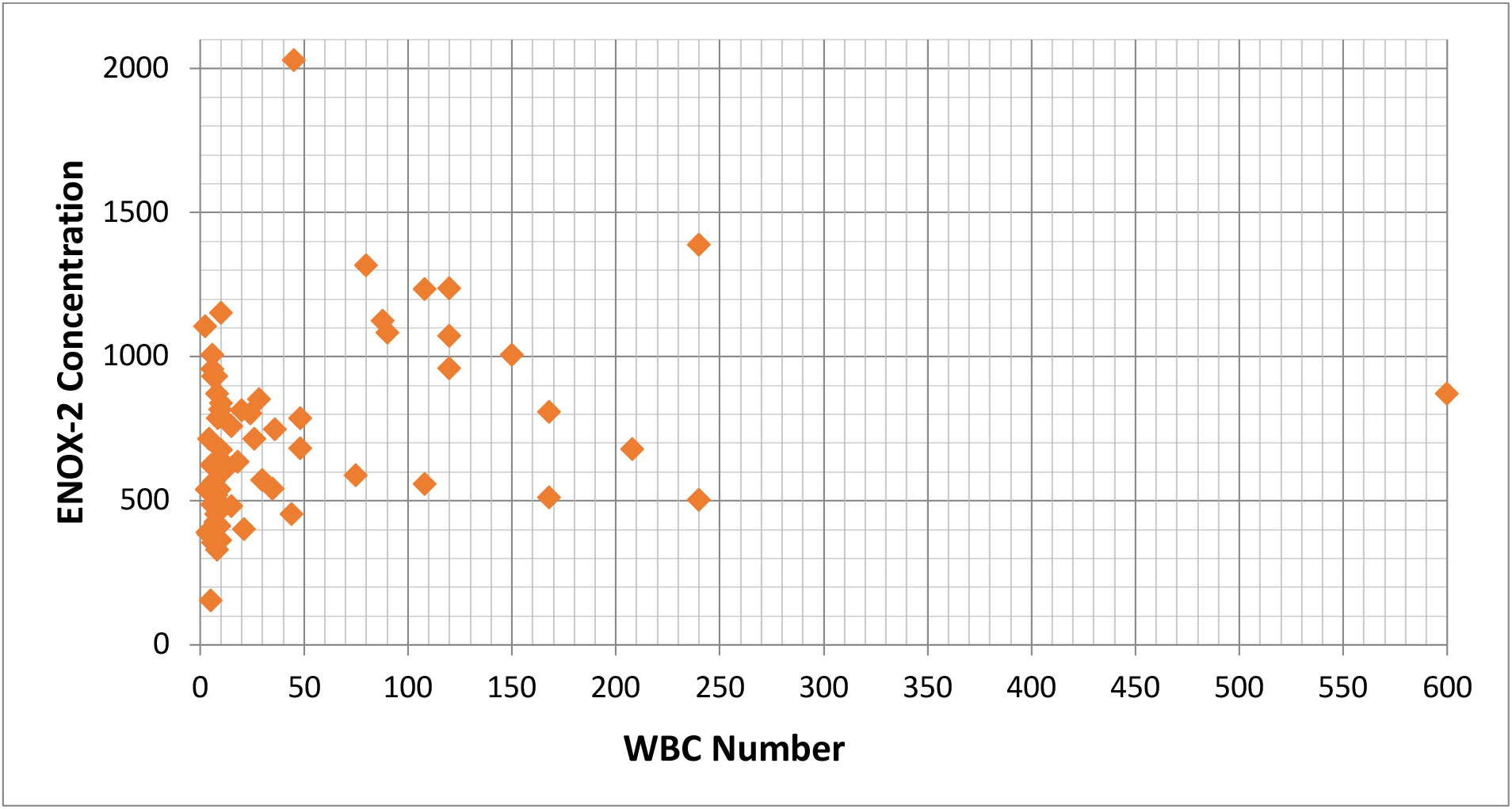
Analysis of a potential relationship between ENOX2 protein levels and white blood cells (WBCs). No correlation was observed between ENOX2 plasma concentrations and WBC count in CML patients at diagnosis.

**S5 Fig:**
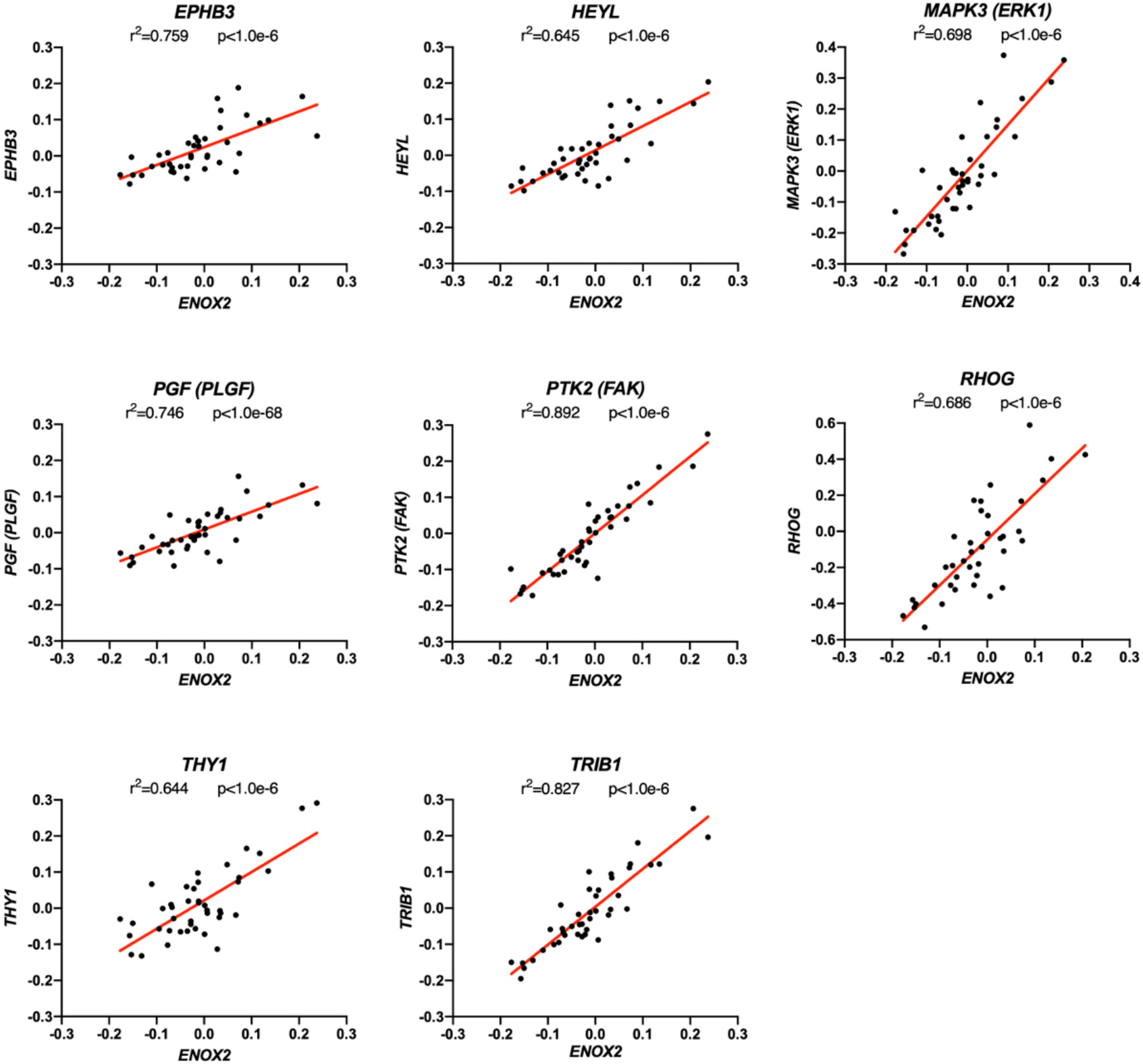
Several potentially relevant protein-coding genes display a mRNA expression significantly correlated to *ENOX2* mRNA expression. Linear regression reveals a high positive correlation degree between *ENOX2* and *EPHB3*, *HEYL*, *ERK1*, *PlGF*, *FAK*, *RHOG*, *THY1*, *TRIB1* gene mRNA expression. The r2 and p parameters are shown on each graph.

**S1 Table:** Patient characteristics (cohort *ENOX2* mRNA expression).

| Patient |  |  | Molecular follow-up<br>(%*BCR-ABL1/ABL1*<sup>IS</sup>) |  | TKI discontinuation |  |  |  |
| --- | --- | --- | --- | --- | --- | --- | --- | --- |
| N° | Sex | Age at diagnosis<br>(years) | Last test<br>(years following<br>diagnosis) | Molecular<br>response | Yes/No | Time to cessation<br>(years following<br>diagnosis) | Result | Duration of TFS<br>(years) |
| 1 | M | 55.2 | 11.9 | MR5 | Yes | 4.1 | TFS | 7.8 |
| 2 | M | 60.1 | 3.0 | MR3 | No |  |  |  |
| 3 | F | 45.2 | 12.6 | MR4.5 | Yes | 6.5 | TFS | 6.1 |
| 4 | M | 76.2 | 7.9 | MR4.5 | No |  |  |  |
| 5 | M | 45.5 | 9.2 | MR4.5 | No |  |  |  |
| 6 | F | 43.7 | 12.0 | MR5 | Yes | 5.6 | TFS | 6.4 |
| 7 | M | 66.9 | 1.8 | MR3 | No |  |  |  |
| 8 | M | 56.5 | 14.6 | MR5 | Yes | 8.1 | TFS | 6.4 |
| 9 | F | 54.3 | 11.3 | MR5 | Yes | 5.7 | TFS | 5.6 |
| 10 | F | 48.6 | 11.1 | MR5 | No |  |  |  |
| 11 | F | 60.4 | 15.0 | MR2 | No |  |  |  |
| 12 | M | 37.4 | 11.3 | MR5 | Yes | 4.1 | Recurrence |  |
| 13 | M | 58.5 | 8.2 | MR4.5 | No |  |  |  |
| 14 | M | 64.2 | 15.4 | MR5 | No |  |  |  |
| 15 | M | 60.5 | 15.3 | MR3 | Yes | 7.5 | Recurrence |  |
| 16 | M | 21.5 | 12.3 | MR4.5 | No |  |  |  |
| 17 | F | 56.8 | 11.3 | MR5 | Yes | 8.0 | Recurrence |  |
| 18 | M | 71.0 | 15.4 | MR5 | Yes | 3.3 | Recurrence |  |
| 19 | F | 34.9 | 17.4 | MR3 | No |  |  |  |
| 20 | F | 84.8 | 7.5 | MR2 | Yes | 6.3 | Recurrence |  |
| 21 | M | 48.6 | 13.3 | MR4.5 | Yes | 7.9 | TFS | 5.3 |
| 22 | F | 57.9 | 5.6 | MR4.5 | No |  |  |  |
| 23 | M | 73.6 | 4.3 | MR3 | No |  |  |  |
| 24 | M | 69.6 | 4.4 | MR4.5 | No |  |  |  |
| 25 | M | 72.3 | 14.7 | MR5 | Yes | 4.5 | Recurrence |  |
| 26 | M | 64.8 | 11.6 | MR4 | No |  |  |  |
| 27 | F | 45.7 | 13.1 | MR5 | Yes | 2.7 | TFS | 10.4 |
| 28 | M | 58.8 | 9.0 | MR3 | No |  |  |  |
| 29 | M | 74.4 | 11.1 | MR4 | No |  |  |  |
| 30 | M | 56.8 | 6.4 | MR4.5 | Yes | 5.0 | Recurrence |  |
| 31 | F | 40.3 | 13.7 | MR5 | No |  |  |  |
| **32** | M | 50.9 | 13.2 | MR4 | No |  |  |  |
| **33** | M | 83.8 | 4.9 | MR4.5 | No |  |  |  |
| **34** | M | 76.6 | 4.0 | MR4 | No |  |  |  |
| **35** | M | 59.3 | 17.0 | MR4 | Yes | 10.7 | Recurrence |  |
| **36** | F | 84.4 | 2.9 | MR3 | No |  |  |  |

TKI, tyrosine kinase inhibitors; TFS, treatment free remission; Molecular recurrence (recurrence in the table) is defined by the loss of MMR (major molecular response) or MR3.
Molecular responses: MR2, %*BCR-ABL1/ABL1*<sup>IS</sup> < 1%; MR3, %*BCR-ABL1/ABL1*<sup>IS</sup> < 0.1%; MR4, %*BCR-ABL1/ABL1*<sup>IS</sup> < 0.01%;
MR4.5, %*BCR-ABL1/ABL1*<sup>IS</sup> < 0.0032% (or 100000 copies > *ABL1* > 32000 copies if undetectable *BCR-ABL1*);
MR5, %*BCR-ABL1/ABL1*<sup>IS</sup> < 0.001% (or *ABL1* > 100000 copies if undetectable *BCR-ABL1*).

**S2 Table:**
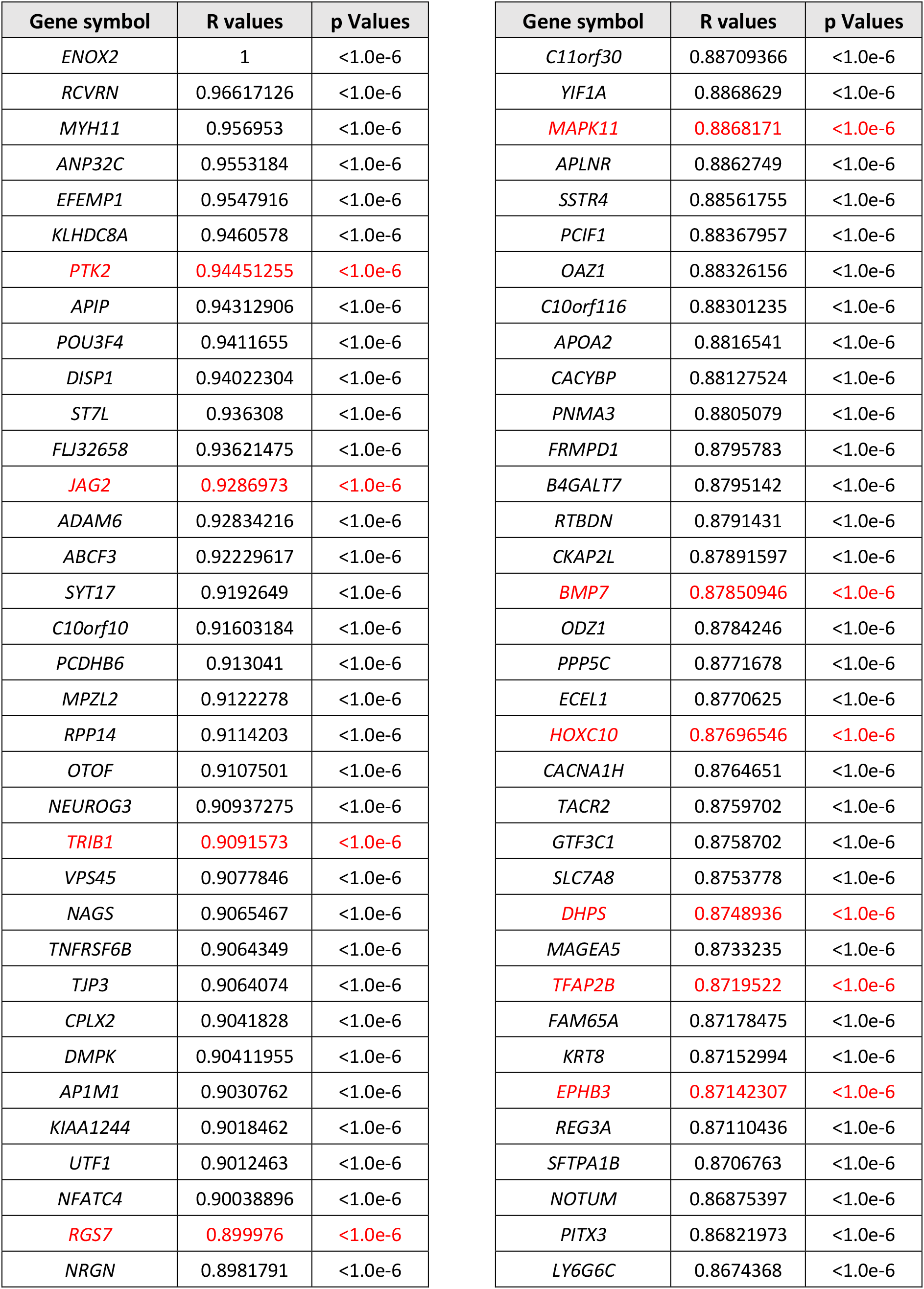
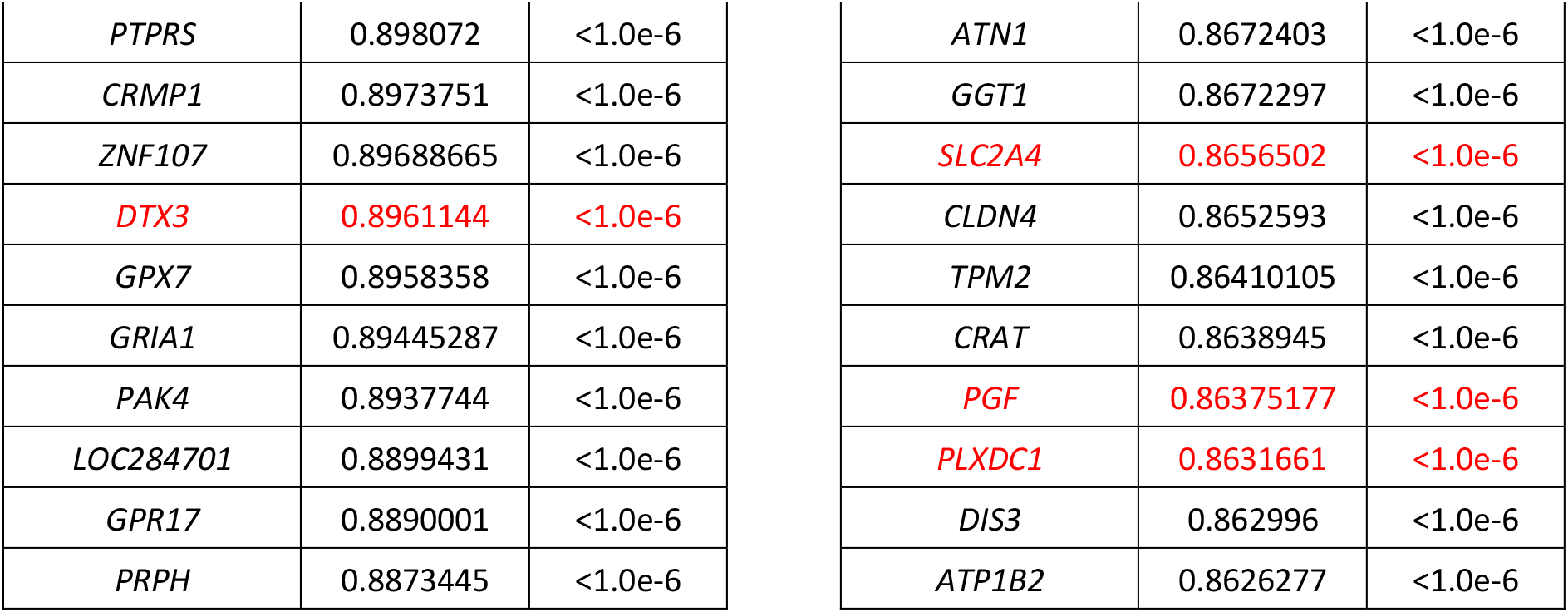
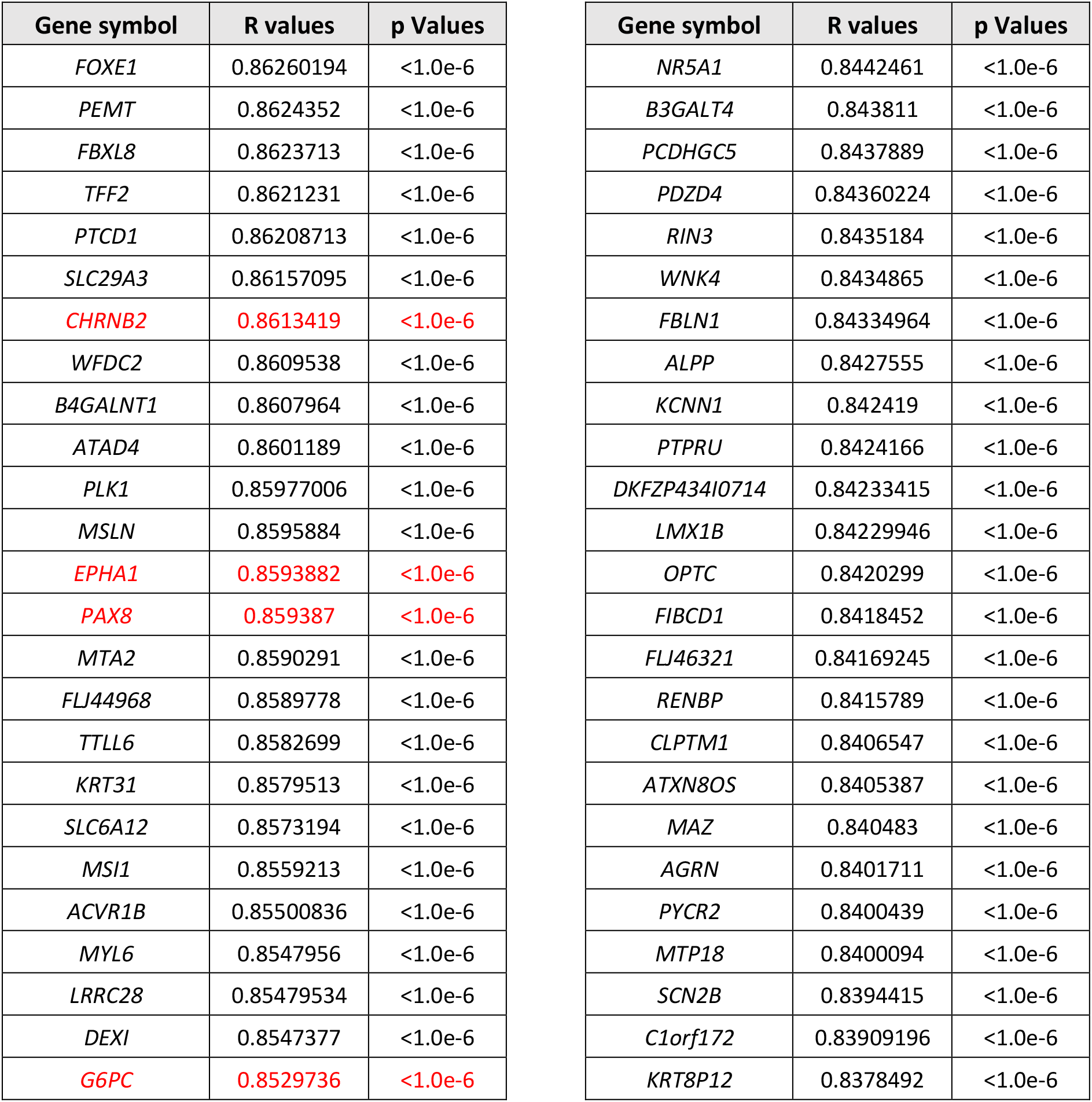
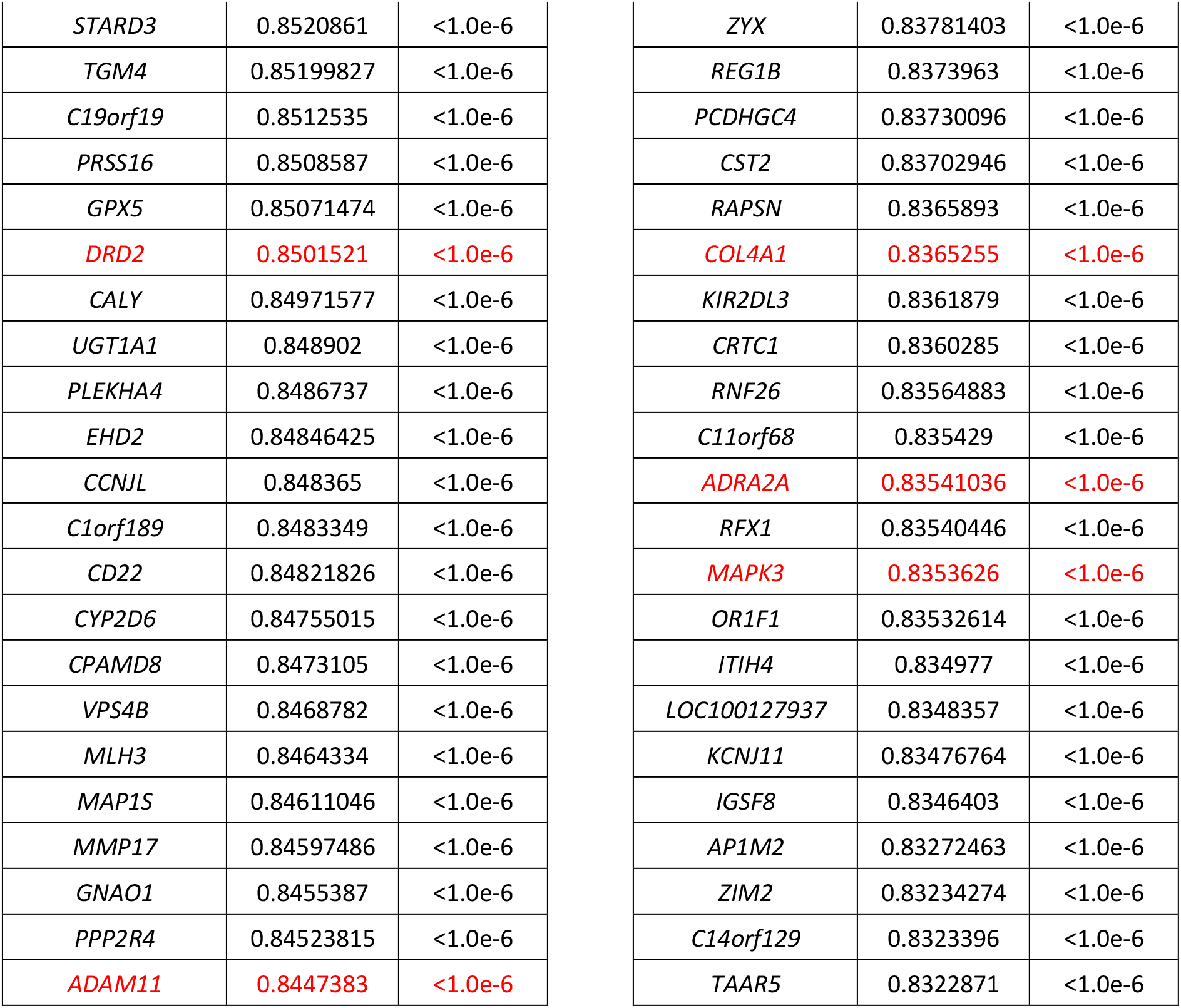
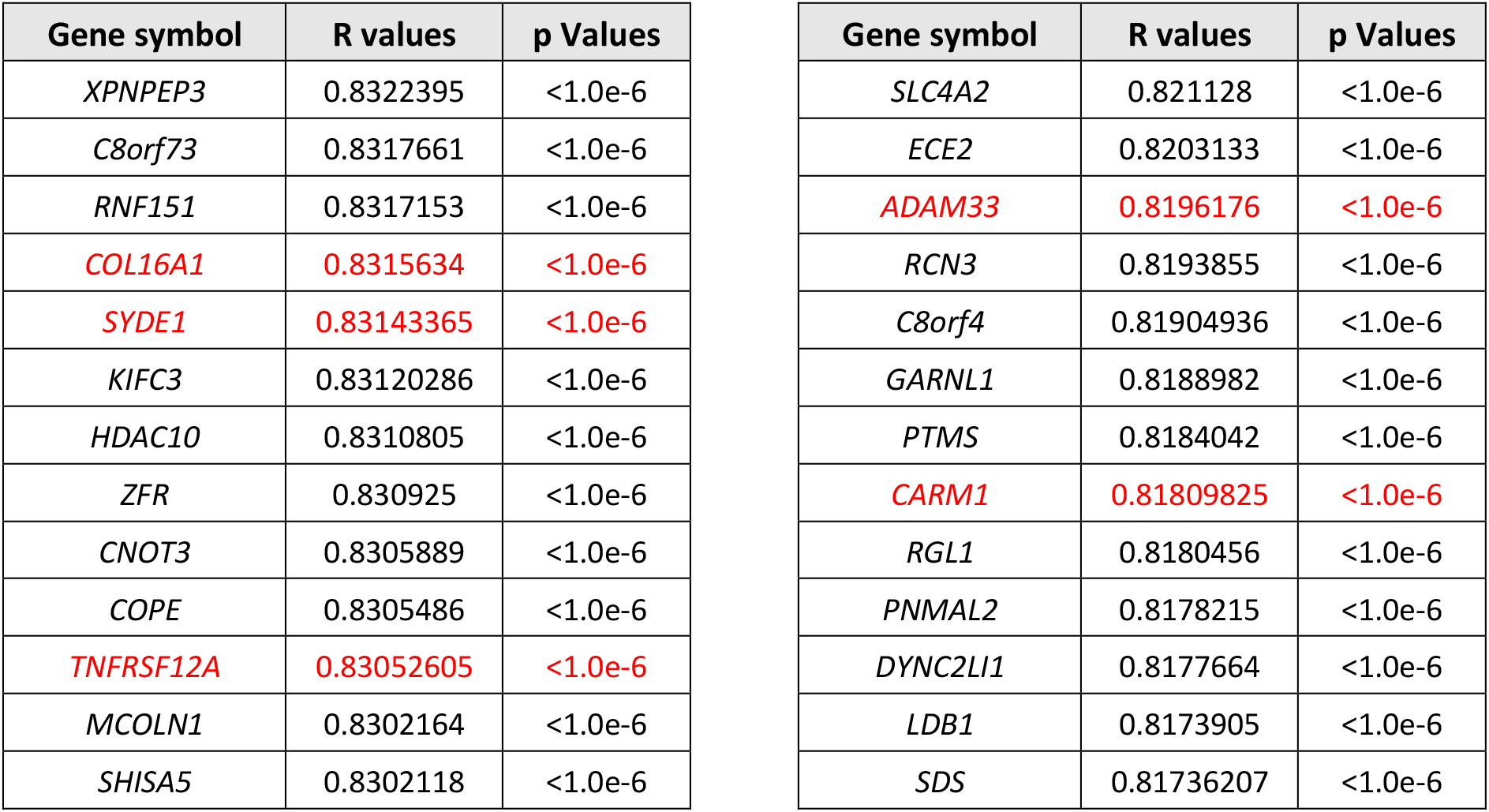
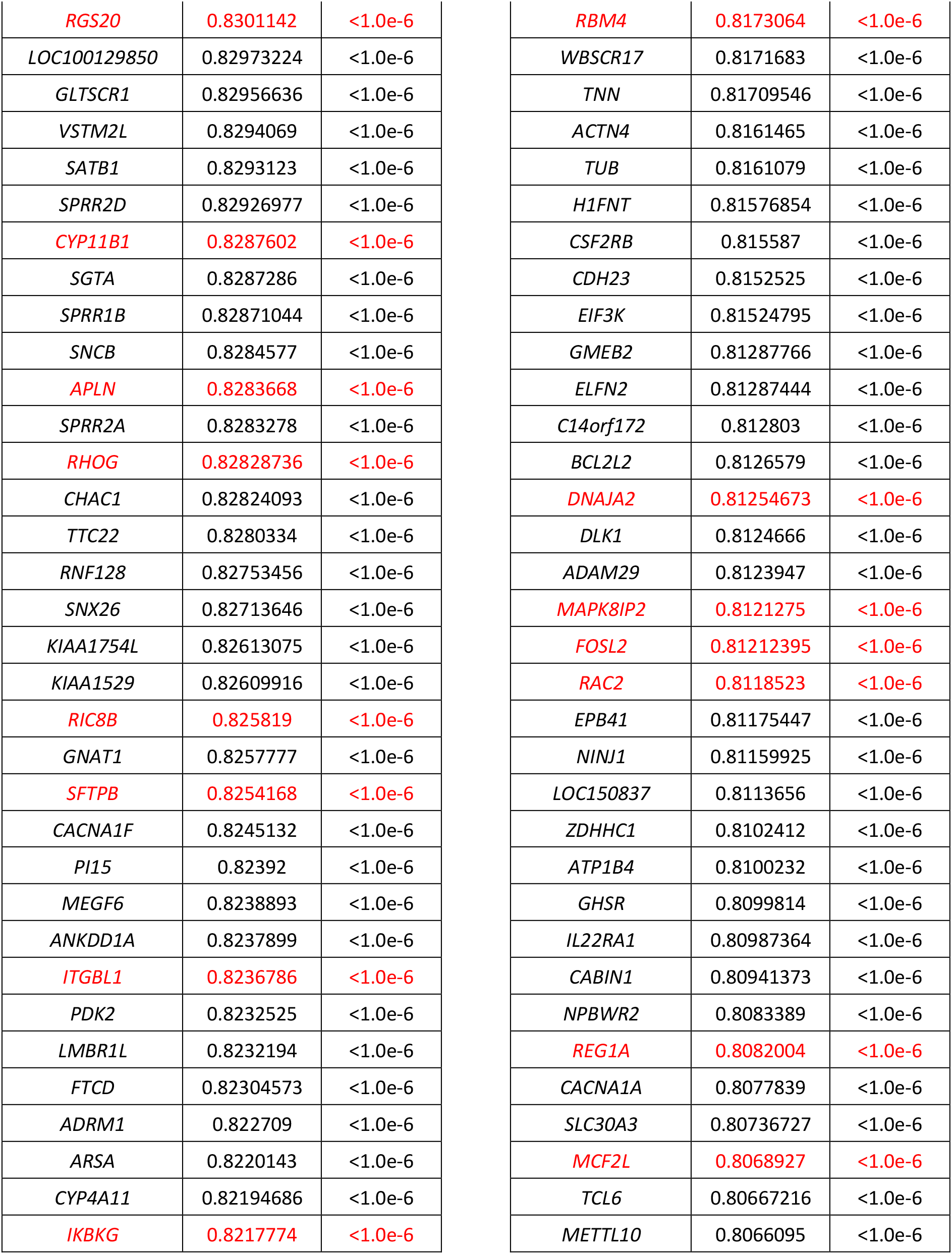
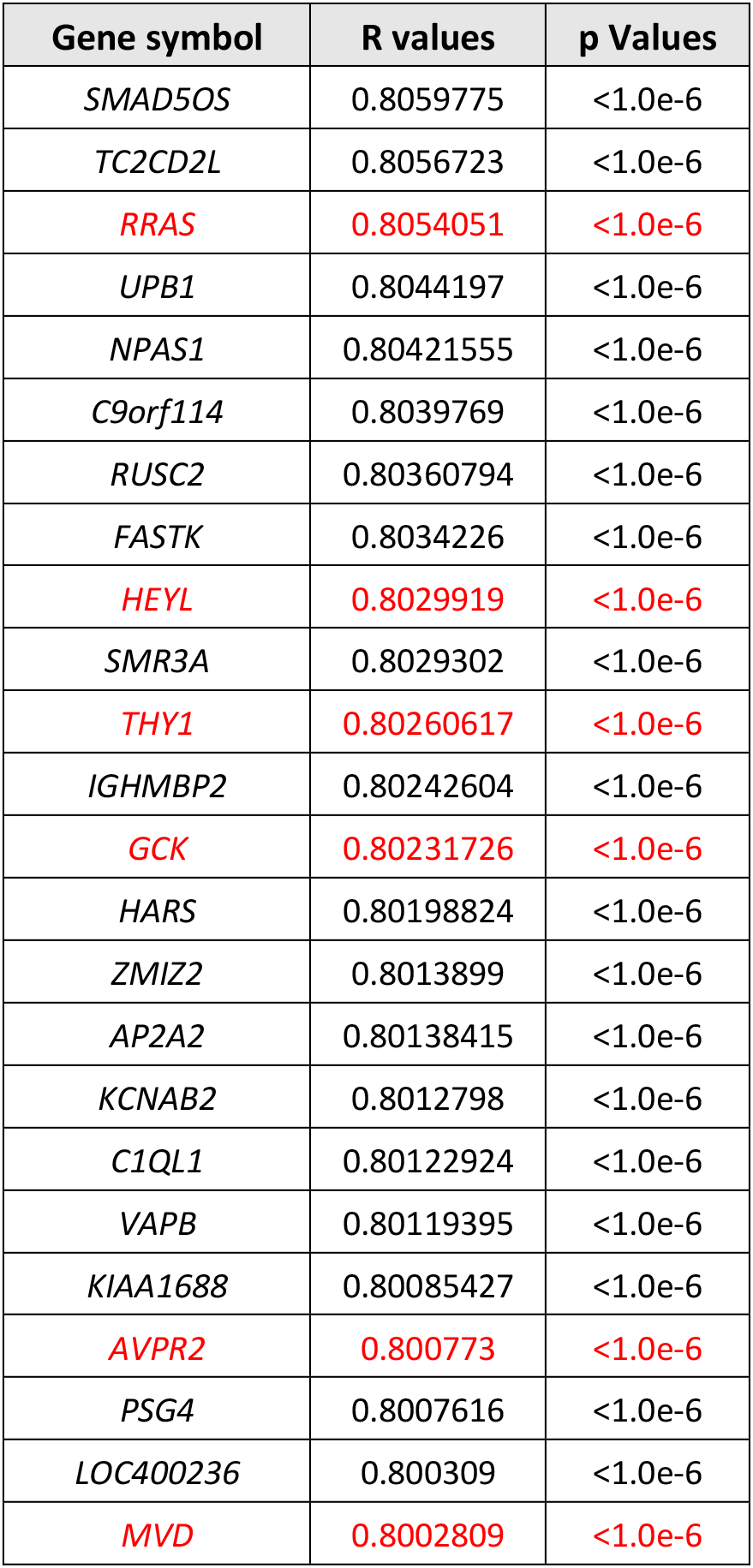
Genes correlated to *ENOX2* mRNA expression. In red, genes known to be involved in essential cell processes.

**S3 Table:**
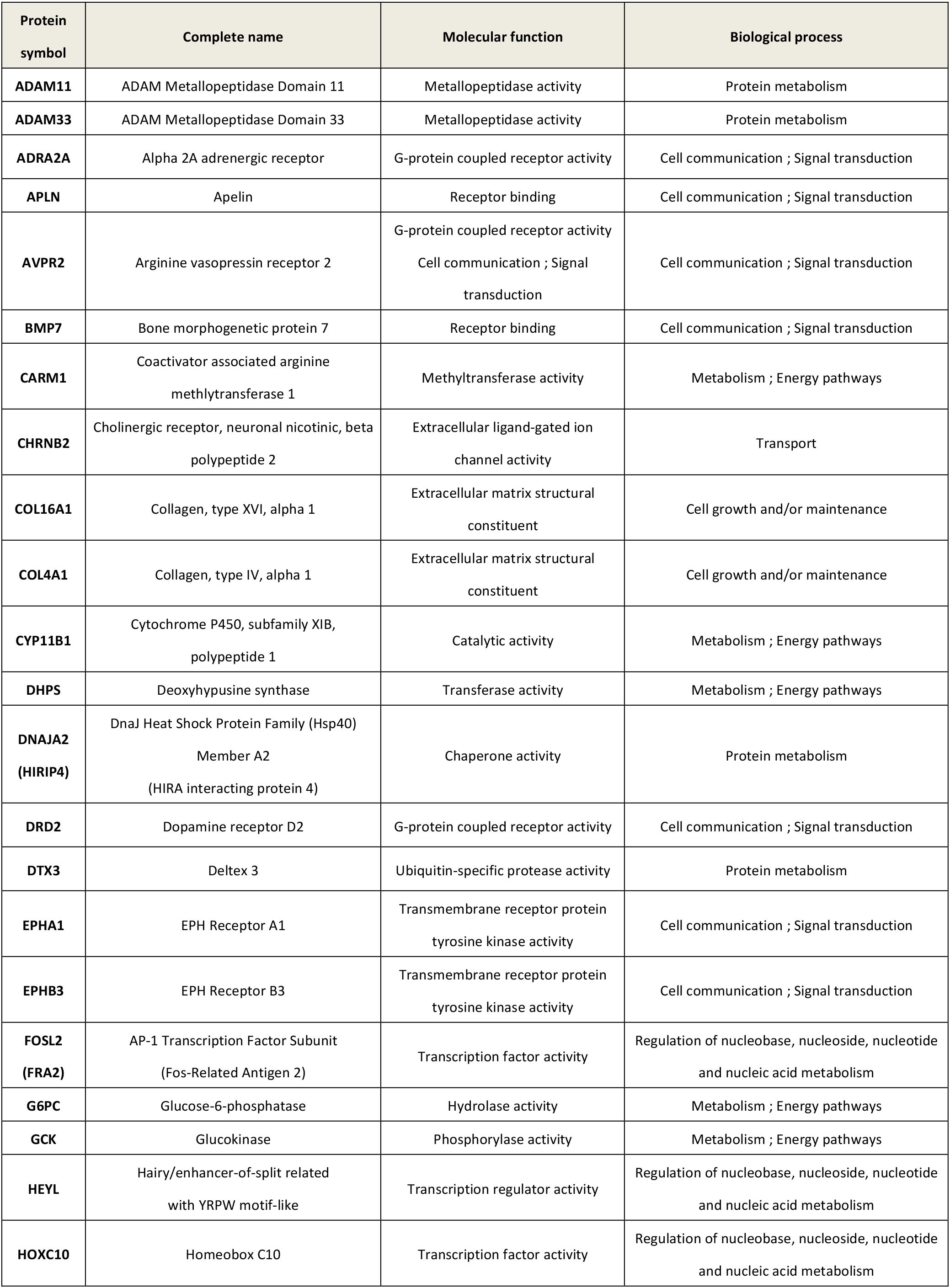
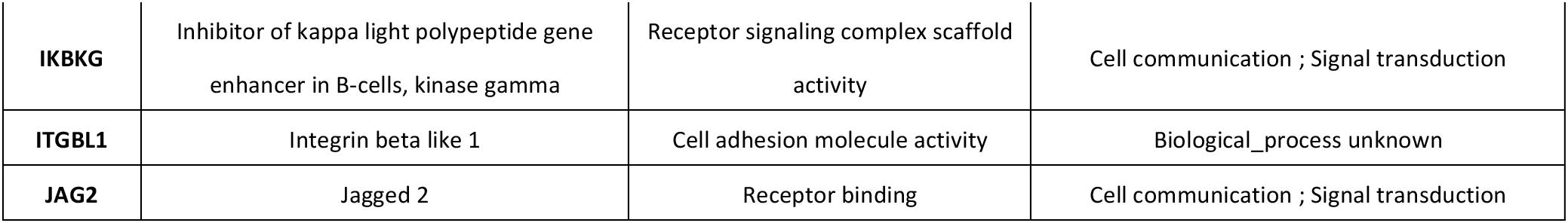
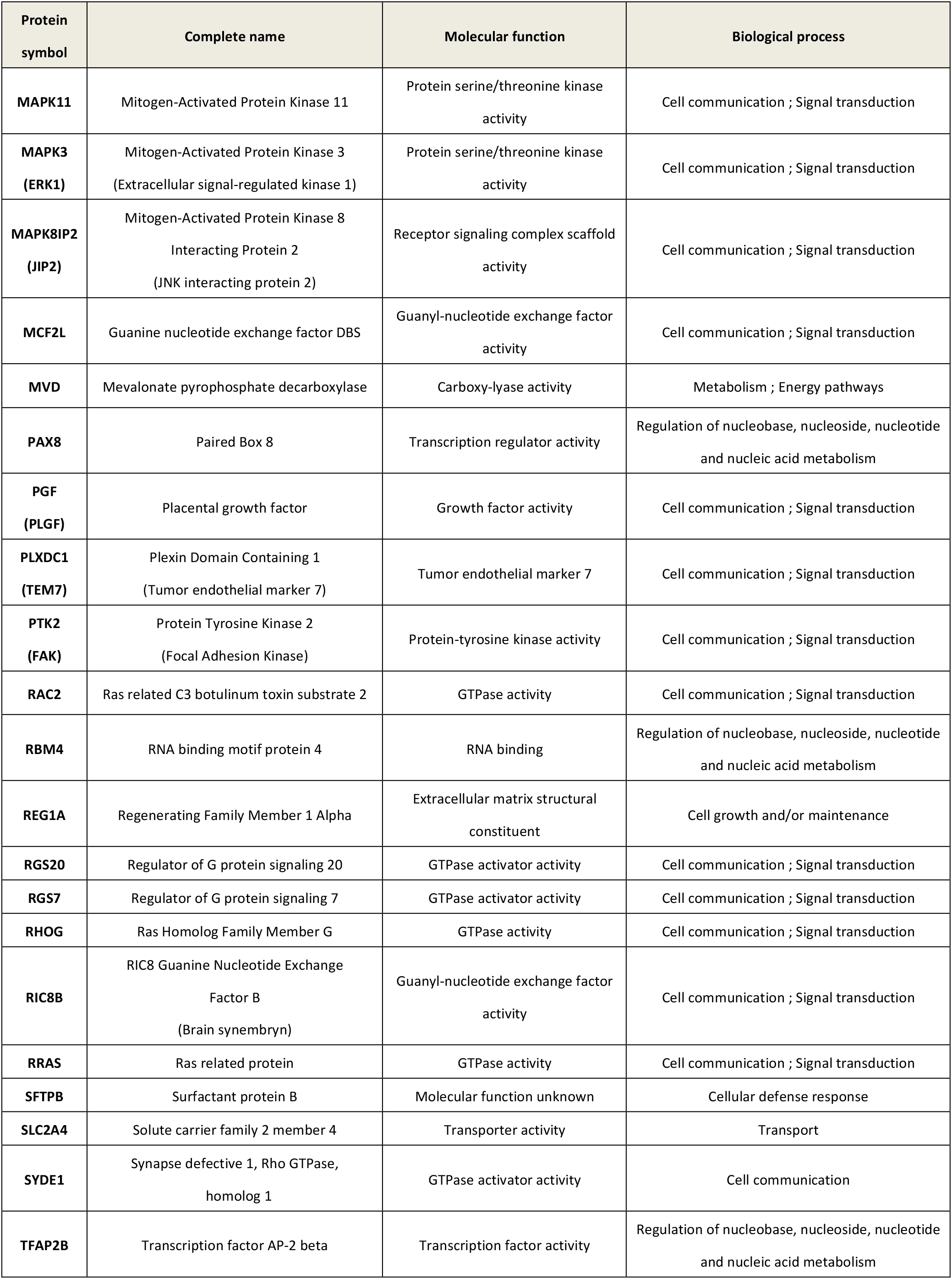
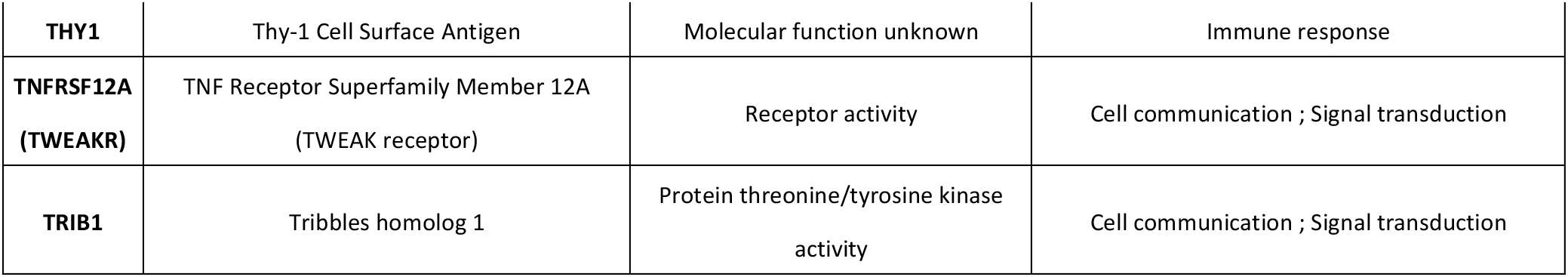
Proteins of ENOX2 potential network involved in critical biological processes.

## Notes

### Competing Interest Statement

The authors have declared no competing interest.

